# Parasitism-mediated horizontal transfer of a functional cytochrome P450 gene entails transposon colonization in newly gained introns

**DOI:** 10.1101/2025.08.25.672084

**Authors:** Eiichiro Ono, Kohki Shimizu, Jun Murata, Tenta Segawa, Akira Shiraishi, Ryusuke Yokoyama, Hiromi Toyonaga, Masaki Takagawa, Manabu Horikawa, Atsushi Hoshino, Koh Aoki

**Author notes:** Corresponding authors: Eiichiro Ono and Koh Aoki. **Author Contributions:** E.O., J.M., and K.A. designed the research. K.S. assembled the genomic DNA sequences of *Cuscuta* species and detected mobile mRNAs. E.O. and H.T. cloned *CYP81Q* genes, constructed expression vectors, performed qRT-PCR, and performed genomic PCR. E.O., J.M., and M.H. performed (+)-sesamin conversion assays. M.H. performed LC-MS analysis. E.O., T.S., M.T., R.Y., and K.A. performed phylogenetic analyses. A.S. performed transcriptome analyses and quantified the ratio of the intron-retained transcripts. R.Y. and M.T. performed similarity searches of the genomic regions flanking *CYP81*-related genes. A.H. and K.A. performed genome synteny and transposon analyses. E.O., T.S., R.Y., and K.A. collected *Cuscuta* samples for (+)-sesamin quantification. E.O., J.M., and K.A. wrote the paper. **Competing Interest Statement:** E.O., H.T., and T.S. are employees of Suntory Global Innovation Center Ltd. All other authors declare no competing financial interests.

## Abstract

Plants produce a wide variety of specialized metabolites that are typically present only in specific lineages. However, some specialized metabolites are found sporadically across distantly related plant species. While the latter cases have been explained as outcomes of convergent evolution, the molecular mechanism behind such metabolic evolution has remained largely elusive. Here, we report that parasitic dodders belonging to the genus *Cuscuta* accumulate sesamin, and that this accumulation may be attributed to the acquisition of enzymatically active homologs of *Sesamum indicum CYP81Q1,* which encodes piperitol/sesamin synthase (PSS). Phylogenetic analysis of *CYP81Q* homologs in *Cuscuta* and *Grammica* subgenera supports a trajectory in which ancestral *Cuscuta* species acquired *CYP81Q* from an ancestral host plant of Lamiales through horizontal gene transfer (HGT), and that the gene has been maintained during the speciation of *Cuscuta*. The evolution of the *CYP81Q* genes was accompanied by sequential intron gains, which likely involved the colonization of transposons. Experiments involving *C. campestris* and *S. indicum* suggested that expression of the host *CYP81Q* gene could be induced by parasitism, and physical connection to the host plant allowed the transfer of genetic elements to *Cuscuta*. These data suggest that parasitism-mediated HGT contributed to the transfer of a gene encoding a key enzyme in specialized lignan metabolism to *Cuscuta*, and that the acquired metabolic gene underwent structural modification while retaining its enzymatic function in dodders.

## Introduction

*Cuscuta* spp. (Convolvulaceae, Solanales), commonly known as dodders (Fig. 1A), are obligate parasitic plants with a broad host range (1-3). These plants are rootless and leafless, and therefore possess limited or no photosynthetic capacity (4). To obtain the sugars and minerals required for survival and proliferation, *Cuscuta* has evolved the ability to form connections with autotrophic host plants through a specialized organ called a haustorium.

**Figure 1.**
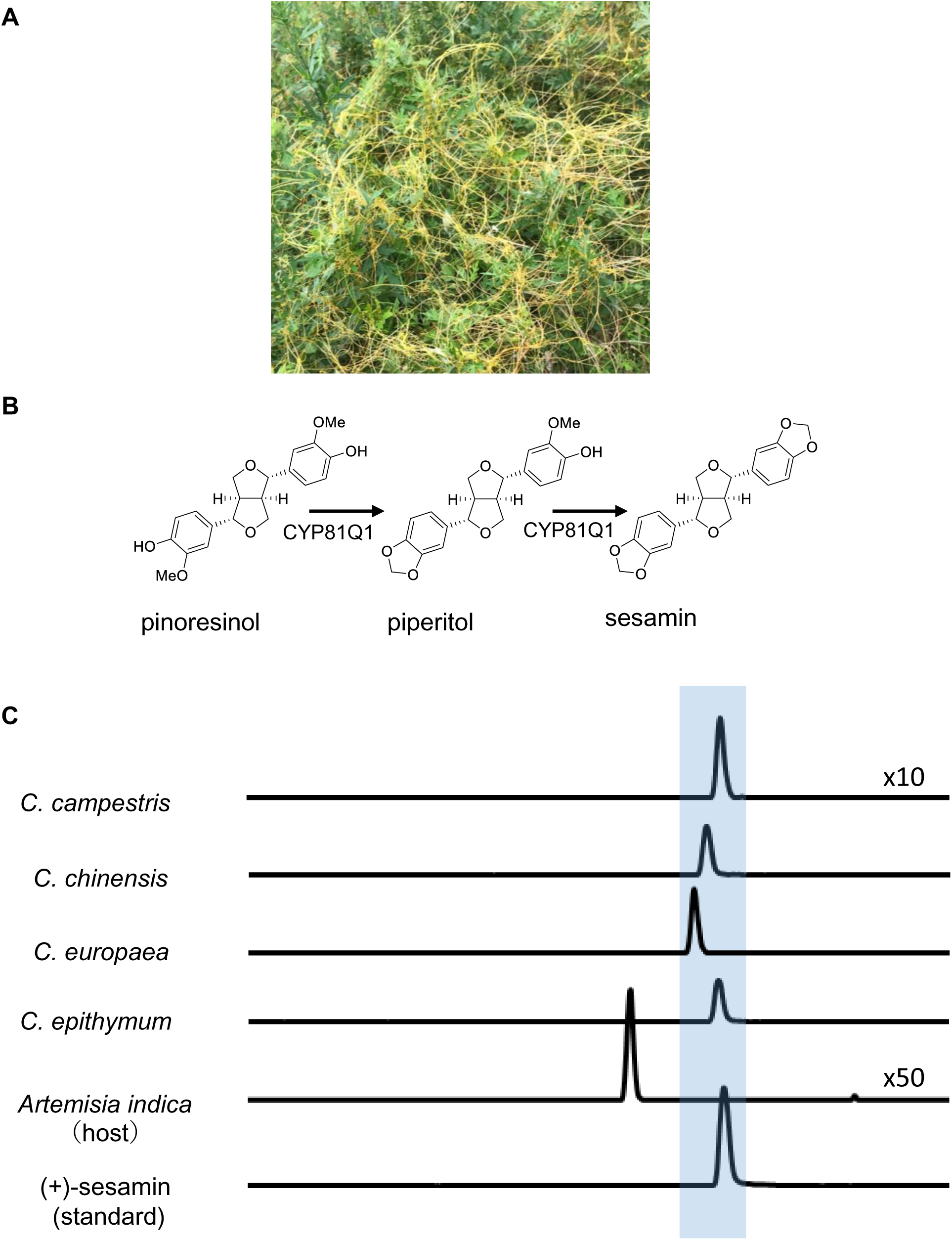
Accumulation of sesamin in *Cuscuta* species: (A) *Cuscuta campestris* parasitizing host plants (*Artemisia indica*) along the coast of Lake Biwa, Shiga Prefecture, Japan. (B) Structure of lignans detected by liquid chromatography-mass spectrometry (LC-MS) analysis. (C) MS chromatograms (m/z 377.10, retention time 20–30 min) of *Cuscuta* extracts (top to bottom): floral clusters containing seeds of *C. campestris*, seeds of *C. chinensis*, seeds of *C. europaea*, stems of *C. epithymum*, leaves of Japanese mugwort (*A. indica*), and an authentic strand of (+)-sesamin. (+)-Sesamin is highlighted with blue marker.

Despite their reduced photosynthetic ability, *Cuscuta* plants are known to produce a variety of bioactive compounds. *Cuscuta* spp. are used in Asian traditional herbal medicine for their purported anti-aging, anti-inflammatory, hepatoprotective, analgesic, and aphrodisiac effects (5). Previous studies have demonstrated that *Cuscuta* spp. contain a variety of specialized bioactive metabolites including flavonoids, steroids, and alkaloids (5, 6), which may represent adaptive metabolic traits associated with parasitism or other aspects of their unique physiology.

Specialized metabolites are generally lineage-specific and restricted to phylogenetically related groups, known as chemotaxonomic groups, due to a shared biosynthetic origin inherited from a common ancestor, e.g., isoflavonoids in Fabaceae. However, certain specialized metabolites are found sporadically in phylogenetically unrelated plants, resulting in metabolic patchiness in chemotaxonomic patterns. For example, sesamin, a specialized lignan, is found in seeds of *Sesamum* spp. (sesame, Pedaliaceae, Lamiales) and in its phylogenetic relatives such as Paulowniaceae, but is also observed in phylogenetically distant plants such as *Piper* spp. (black pepper), *Magnolia* spp. (a basal angiosperm), and *Ginkgo* (a gymnosperm) (7–9) (*SI Appendix*, Fig. S1). In *S. indicum*, a cytochrome P450 monooxygenase, SiCYP81Q1, also known as piperitol/sesamin synthase (PSS), catalyzes the formation of two methylenedioxy bridges (MDBs) on the two aromatic rings through sequential conversion of pinoresinol to piperitol and then to sesamin (10) (Fig. 1B). Recent studies indicate that accumulation of sesamin and its related metabolites in phylogenetic relatives of *S. indicum* coincides with the presence of functional *CYP81Q* genes that exhibit high sequence similarity to *SiCYP81Q1* (10-12). Notably, *Cuscuta* is, to date, the only genus in the Convolvulaceae family known to accumulate sesamin (7, 13-15).

The presence of sesamin in *C. palaestina,* which parasitizes host plants that do not accumulate sesamin, suggests that *C. palaestina* possesses the biosynthetic enzyme required for sesamin production *in planta,* rather than acquiring sesamin from its host (15). Since pinoresinol, the precursor of sesamin, is widespread across land plants, investigation of sesamin biosynthesis in *Cuscuta* spp. provides a model for understanding the molecular mechanisms that underlie the sporadic distribution of specialized metabolites.

Given that the PSS genes in Lamiales are currently the only genes known to encode proteins with sesamin synthase activity, at least three possible explanations exist for the sporadic occurrence of sesamin in *Cuscuta* spp.: 1) functional differentiation or gene loss, 2) convergent evolution of a sesamin synthase, and 3) horizontal gene transfer (HGT) of a sesamin synthase gene. A previous report on the functional differentiation of sesamin synthase showed that *S. alatum* CYP81Q3 produces pluviatilol instead of sesamin, resulting in the exceptional absence of sesamin in *S. alatum* among the *Sesamum* spp. (12). Functional differentiation or gene loss is likely restricted to relatively small taxonomic groups, as it depends on preceding metabolic radiation coupled with a common biosynthetic gene. Conversely, convergent evolution can occur across distant lineages, independently of metabolic radiation. As a result, convergently evolved genes typically share low sequence similarity due to their distinct genetic origins. HGT provides an alternative explanation for the presence of sporadic metabolites. Recent studies have shown that *Cuscuta* plants can acquire genes from host plants (3,16–18). Therefore, HGT is a plausible mechanism in parasitic plants such as *Cuscuta*, *Striga*, and *Orobanche* spp., given that the intimate physical interactions formed with host plants provide opportunities for gene transfer (16, 17). A characteristic of HGT-derived genes is that they are structurally similar to homologs in host plants. For example, *STRICTOSIDINE SYNTHASE-LIKE* genes in *C. australis* exhibit much higher sequence similarity to orthologs in Brassicaceae than with genes in *Cuscuta*’s close relatives, suggesting that HGT from Brassicaceae host plants to the parasite might have occurred (18). Although these findings support the involvement of HGT, the biochemical functionality and biological impact of such HGT-acquired genes have yet to be elucidated.

In this study, we attempted to clarify the molecular basis underlying the occurrence of sesamin and its structurally related specialized lignans bearing MDBs in *Cuscuta* species. We identified homologs of *SiCYP81Q1* in *Cuscuta* spp., all of which are functionally active as PSSs. Comparative analysis of genomic regions adjacent to *CYP81Q* genes revealed that *Cuscuta CYP81Q* genes are integrated into genomic regions exhibiting conserved synteny among *Cuscuta* species and with the close relative *Ipomoea*, suggesting that *CYP81Q* was acquired by HGT. The presence of *CYP81Q* supports an evolutionary scenario in which a common ancestor of the *Cuscuta* and *Grammica* subgenera acquired a functional *PSS* gene from an ancestral host plant capable of producing sesamin, likely within Lamiales, through parasitism-mediated HGT (pHGT). We propose that pHGT represents the metabolic origin of sesamin in non-Lamiales plants. Notably, intron gain was preferentially observed in these horizontally transferred genes, including *CYP81Q* genes. This genomic signature may reflect transposon colonization of *trans*-genes in dodders.

## Results

### Identification of Cuscuta CYP81Q-related genes

The presence of sesamin and structurally related lignans in seeds of *Cuscuta chinensis*, *C. australis*, and *C. palaestina* (13–15), all within the subgenus *Grammica*, led us to examine whether other *Cuscuta* plants also contain sesamin. We analyzed lignans in seeds of *C. campestris* and *C. chinensis* parasitizing the experimental host plants *Nicotiana tabacum* and *Arabidopsis thaliana*, respectively, which do not produce sesamin, and detected sesamin and related lignans (Fig. 1C, *SI Appendix*, Fig. S2). Moreover, we detected sesamin in both *C. europaea* and *C. epithymum* of the subgenus *Cuscuta*, but not in a natural host plant of *C. campestris*, *Artemisia indica* (Fig. 1A). These results indicate that sesamin is synthesized *de novo* in *Cuscuta* plants and is not acquired from host plants.

To elucidate the molecular basis of sesamin biosynthesis in *Cuscuta* species, we investigated genome and transcriptome data from multiple *Cuscuta* species (assemblies: GCA_003260385.1, GCA_900332095.2, GCA_945859875.1, GCA_945859915.1) (3, 19) using the Basic Local Alignment Search Tool (BLAST) (20) with *SiCYP81Q1* (AB194714, NM_001319691) as the query. We identified *CYP81Q1* homologs in species belonging to subgenera *Grammica* and *Cuscuta*: *CausCYP81Q111* (C065N002E0.1, RAL50776.1) in *C. australis*, *CcamCYP81Q110* (Cc046292, VFQ70270.1) and *CcamCYP81Q111* (Cc015414, VFR00937.1) in *C. campestris*, *CchiCYP81Q-1* (LC829580) and *CchiCYP81Q-2* (LC829581) in *C. chinensis*, *CeurCYP81Q* (CEURO_LOCUS2477, CAH9067102.1) in *C. europaea*, *CepiCYP81Q-1* (CEPIT_LOCUS12958, CAH9094665.1) and *CepiCYP81Q-2* (CEPIT_LOCUS39506, CAH9141928.1) in *C. epithymum* (Fig. 2). The putative amino acid sequences of these genes share 78–95% identity with SiCYP81Q1 (*SI Appendix*, Data S1).

**Figure 2.**
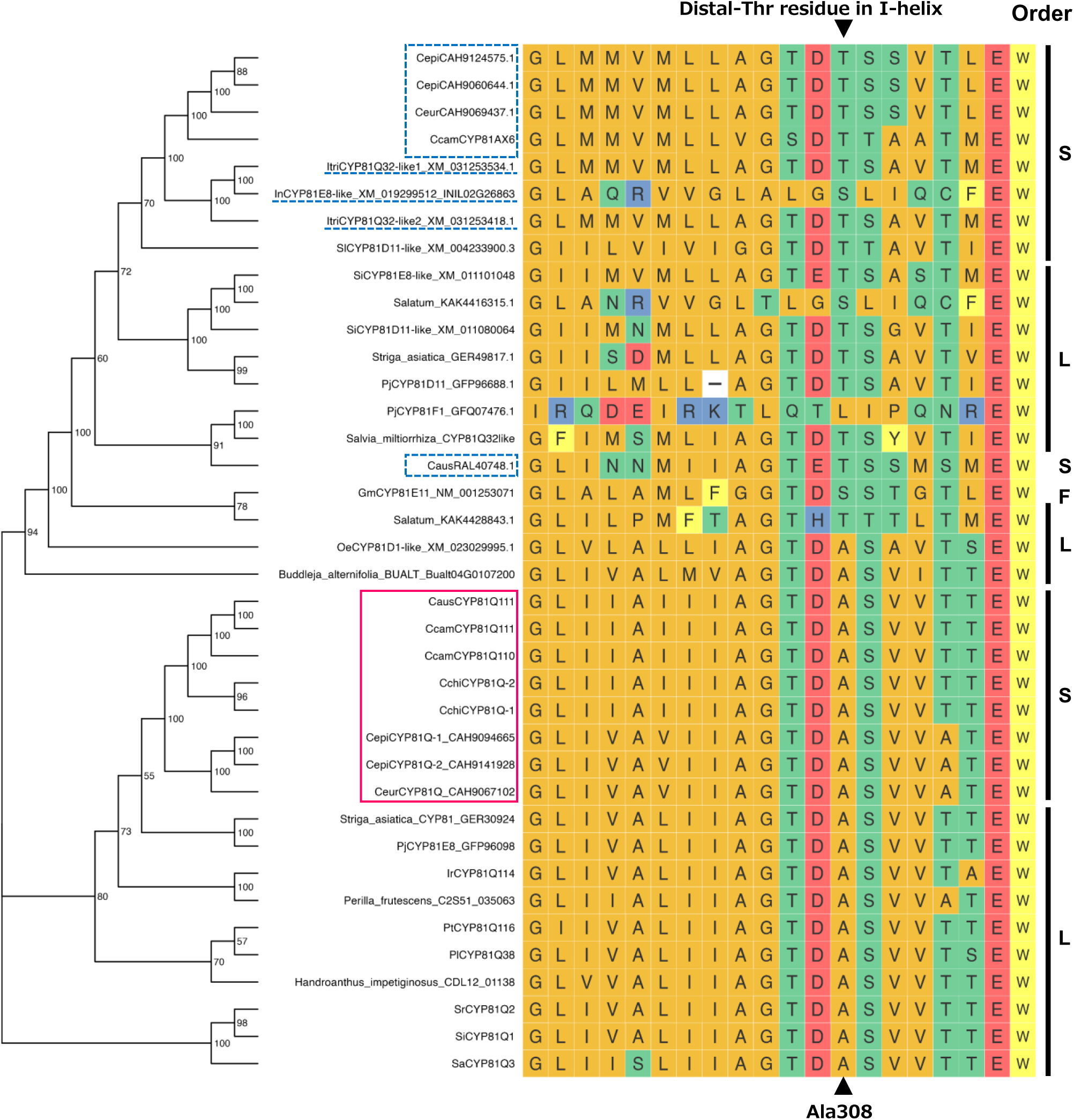
Phylogeny of *CYP81Q*-related genes: (Left) Maximum likelihood phylogenetic tree of *CYP81Q* homologs and related genes in plants belonging to order Solanales and Lamiales. Protein IDs are listed in *SI Appendix*, Data S2. Numbers on branches indicate bootstrap values. Red rectangle: *Cuscuta* CYP81Q homologs. Blue rectangle: *Cuscuta* CYP81 homologs from different subfamilies. Blue dotted line: *Ipomoea* proteins with highest similarity to SiCYP81Q1 in each species. (Center) Alignment of amino acid sequences near the distal-Thr residue in the I-helix. Triangles indicate the position of threonine (Thr) or substituted alanine (Ala). (Right) Taxonomic orders of each plant species: F, Fabales; L, Lamiales; S, Solanales.

We also identified partial sequences of *CYP81Q1* homologs in *Cuscuta* species for which whole-genome sequences are not yet available, and performed phylogenetic analysis using the exon 2 sequences from *C. americana*, *C. californica*, and *C. gronovii*. Phylogenetic analysis of these exon 2 sequences, together with those from the eight sequenced species, was broadly consistent with the phylogeny of *Cuscuta* spp., suggesting a monophyletic origin of these genes within dodder speciation (*SI Appendix*, Data S2, Fig. S3).

Furthermore, we identified *CcamCYP81AX6* (Cc047366, VFQ75312.1) as a third homolog of *SiCYP81Q1* in *C. campestris*, as well as homologs of *CcamCYP81AX6* from *C. australis*, *C. europaea*, and *C. epithymum*. However, CcamCYP81AX6 shares a relatively low amino acid identity with SiCYP81Q1 (51%). Phylogenetic analysis showed that *CcamCYP81AX6* homologs form a distinct phylogenetic cluster together with *CYP81D/E*-related genes from *I. nil*, *I. triloba*, and several Lamiales species, with a single homolog from *C. australis* (CausRAL40748.1) being an exception (Fig. 2, *SI Appendix*, Data S3, Data S4). This cluster is distinct from that harboring *Cuscuta CYP81Qs* (Fig. 2).

Notably, the amino acid sequence of SiCYP81Q1 has a unique Ala (Ala308) residue in the distal I-helix that is crucial for the catalysis of MDB formation. Typically, cytochrome P450s have a conserved Thr residue (distal-Thr) in this position (Fig. 2, *SI Appendix*, Data S4) (11, 21). *SiCYP81Q1* homologs in *C. australis*, *C. campestris*, *C. chinensis*, *C europaea*, and *C. epithymum* all possess an Ala residue at the site corresponding to the distal-Thr residue (Fig. 2). This conservation of a crucial residue for catalysis satisfies the structural criteria for putative MDB-forming enzymes and suggests that these homologs likely catalyze sesamin biosynthesis. In contrast, CcamCYP81AX6-related proteins have Thr at the corresponding position.

### Identification of CYP81Q-related genes in Lamiales and Solanales

To clarify the distribution of *CYP81Q*-related genes, we investigated whether other species within the Lamiales and Solanales possess *CYP81Q* homologs (Fig. 2, *SI Appendix*, Data S3, Data S4). Species belonging to Lamiales, such as *Buddleja alternifolia*, *Phryma leptostachya*, *Paulownia tomentosa*, *Phtheirospermum japonicum*, *Striga asiatica*, *Isodon rubescens*, and *Perilla frutescens*, all have *CYP81Q* homologs. CYP81Q proteins of these Lamiales species have an Ala residue at the site corresponding to the distal-Thr residue (Fig. 2). *CYP81Q* homologs of *Cuscuta* species are embedded within the same clade as these Lamiales *CYP81Q* genes, including those from *P. leptostachya, P. tomentosa*, *I. rubescens*, *P. japonicum*, and *S. asiatica*, all of which retain alanine at the site corresponding to the distal-Thr residue.

On the other hand, although *Ipomoea* is considered to be the closest relative of *Cuscuta*, no *CYP81Q* homologs were identified in the genomes of *Ipomoea* species. *Ipomoea triloba* (assembly: GCA_003576645.1) (22) contains two genes, XM_031253481 and XM_031253534, and *I. nil* (assembly: GCA_001879475.1) (23) contains two genes, XM_019299512 and XM_019299461, that show moderate sequence similarity to *SiCYP81Q*1. However, none of these proteins encode a protein with an Ala at the site corresponding to the distal-Thr residue. Similarly, *Solanum lycopersicum* (assembly: GCA_000188115.4) (24) has one gene, XM_004233900, with high similarity to *SiCYP81Q1*, but also retains a Thr at the corresponding site. Phylogenetically, these genes were clustered with *CcamCYP81AX6* homologs from *Cuscuta* spp.

### Comparison of gene synteny in genomic regions containing CYP81Q genes

We next compared gene synteny in the genomic regions containing *CYP81Q* genes across *S. indicum, Cuscuta* species, and related genera in Solanales, including *Solanum* and *Ipomoea*. The analyzed regions ranged in size from 260 kb to 4.2 Mb (Fig. 3, *SI Appendix*, Data S5). In the *S. indicum* genome, *SiCYP81Q1* is located on chromosome 9 (NC_026159.1) (25), between a gene encoding FLOWERING LOCUS C expresser (FLX)-like protein (XP_020554790.1) and a gene encoding an MYB44-like transcription factor (XP_011099313.1). By contrast, *CYP81Q* genes of *Cuscuta* species are located near genes encoding a C3H zinc finger transcription factor, a B3 domain-containing protein, two uncharacterized proteins, an E3 ubiquitin-protein ligase, a methionyl-tRNA formyltransferase, a nucleolin-like protein, and a protease Do-like 7 (gene labels 17 to 25, Fig. 3, *SI Appendix*, Data S5). The order of genes encoding the E3 ubiquitin-protein ligase, uncharacterized protein, methionyl-tRNA formyltransferase (gene labels 20, 22, and 23, Fig. 3) was inverted between the subgenera *Cuscuta* and *Grammica*. In flanking regions upstream and downstream, the genomic order of 16 genes (gene labels 1–12, 14, and 16) was conserved among all *Cuscuta* species. In the genomes of *Ipomoea nil, I. triloba*, and *Solanum lycopersicum*, genes having high similarity to the abovementioned genes (gene labels 1–25) were clustered on the same scaffolds, and their sequential order was conserved. In contrast, no genes similar to those labeled 1–25 were found near *SiCYP81Q1* in *S. indicum*. These findings highlight the irregular presence of *CYP81Q* genes in *Cuscuta* species within *Solanales*.

**Figure 3.**
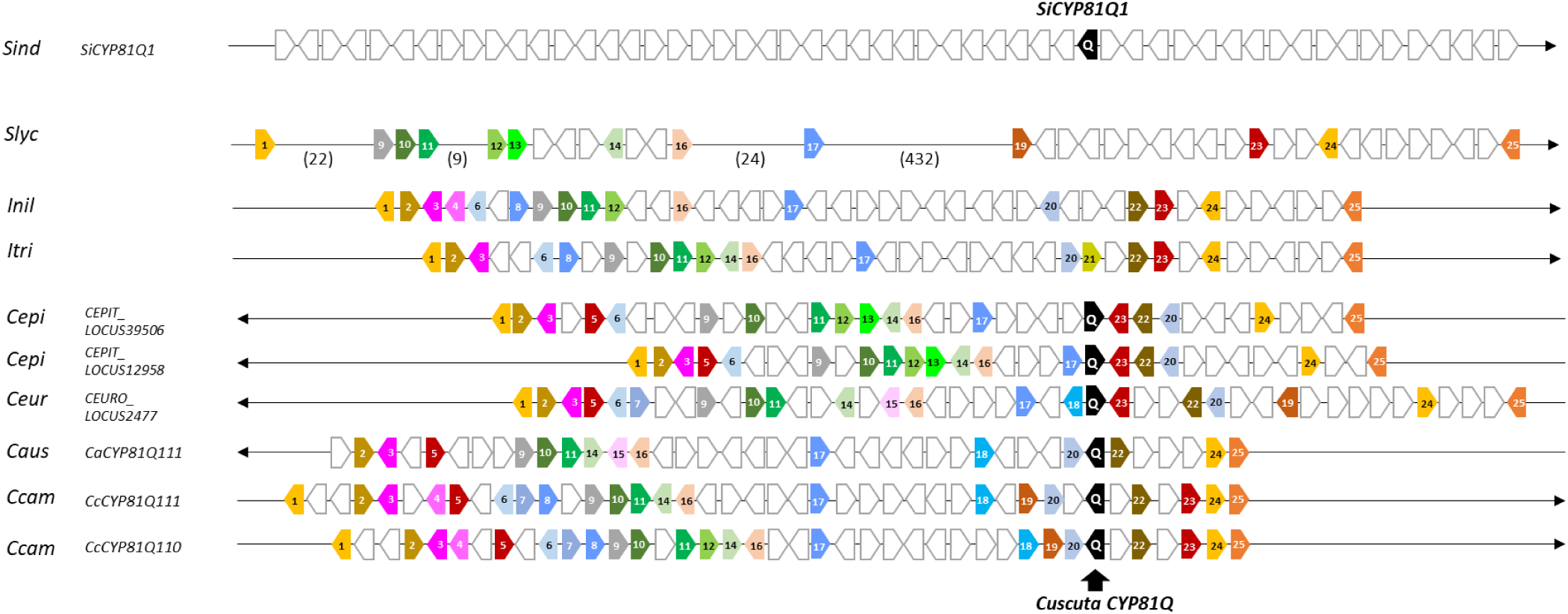
Comparative genomic synteny among *Cuscuta* and related species: Gene synteny in the genomic region surrounding *CYP81Q*-related genes is highly conserved among Solanales species, including *Cuscuta*, *Ipomoea*, and *Solanum*, whereas *Paulownia fortunei* and *Sesamum indicum* (Lamiales) show a distinct genomic structure. *Pfor*, *P. fortune; Sind*, *S. indicum; Slyc*, *Solanum lycopersicum; Inil*, *Ipomoea nil*; *Itri*, *I. triloba*; *Cepi, C. epithymum*; *Ceur*, *C. europaea*; *Caus*, *C. australis*; *Ccam, C. campestris*. Black boxes labeled “Q” indicate *CYP81Q*-related genes. IDs and descriptions of numbered genes are listed in *SI Appendix,* Data S5. Numbers in parentheses indicate the number of genes within each interval. Grey triangles between *Pfor* and *Sind* denote chromosomal inversions.

The discrepancy between the linear phylogenetic relationship and genomic synteny of *Cuscuta CYP81Q* genes suggests that *Cuscuta CYP81Q* genes were horizontally transferred from an ancestral Lamiales species into a genomic region where gene synteny is conserved within Solanales, rather than having gradually differentiated from a Solanales ancestor. Based on synonymous substitution rates, this HGT event is estimated to have occurred between approximately 50.38 and 32.05 million years ago (Ma), corresponding to the inferred divergence time between parasitic *Cuscuta* and non-parasitic *Ipomoea* spp. within Solanales. In contrast, the divergence between *Cuscuta* and *Sesamum* (Lamiales) is estimated at approximately 101.63 Ma *(SI Appendix*, Data S6, Fig. S4).

### Functional evaluation of Cuscuta CYP81Q genes

The co-occurrence of *CYP81Q*-related genes and sesamin in *Cuscuta* species suggests a biochemical role for these genes as PSSs. To evaluate the enzymatic activity of proteins encoded by *Cuscuta CYP81Q* genes, recombinant proteins were co-expressed in a yeast system with *C. campestris* cytochrome P450 reductase, CcamCPR1 (Cc043955, VFQ62263.1) (26). When fed with (+)-pinoresinol, CausCYP81Q111, CcamCYP81Q110, CcamCYP81Q111, or CchiCYP81Q-1 each catalyzed the formation of two MDBs on the two aromatic rings of the (+)-pinoresinol, producing (+)-sesamin via (+)-piperitol (Fig. 4). The stereochemistry of the resulting (+)-sesamin was identical to that generated by SiCYP81Q1 using (+)-pinoresinol as a substrate. In contrast, CcamCYP81AX6, did not exhibit MDB-forming activity for any of the pinoresinol isomers tested (*SI Appendix*, Fig. S5). These results demonstrate that the eight *CYP81Q* genes identified in the genus *Cuscuta* encode functional PSS enzymes and provide a molecular explanation for the presence of sesamin and derived lignans such as cuscutoside in *Cuscuta* species.

**Figure 4.**
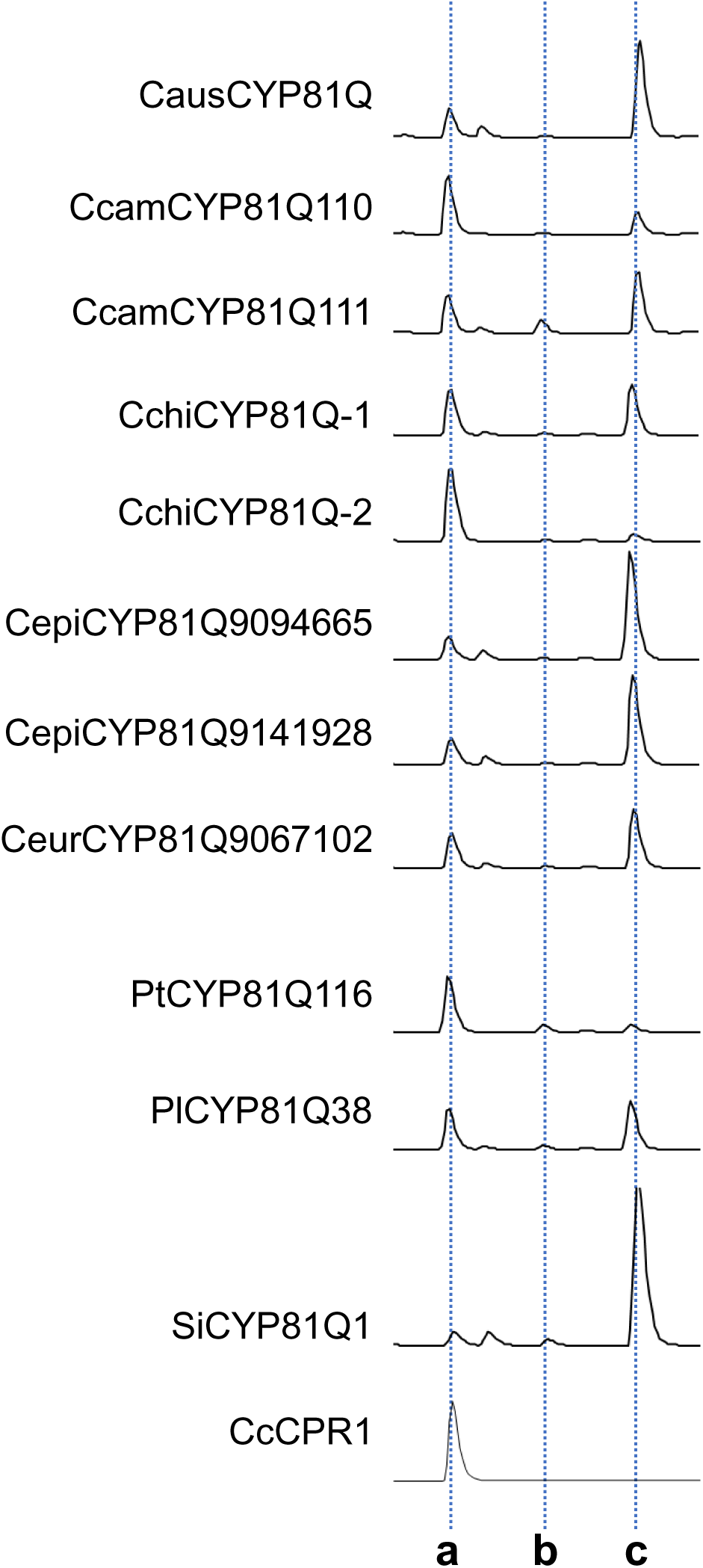
Enzymatic activity of *Cuscuta* PSS protein: Products of yeast enzyme assays analyzed by HPLC. Chromatograms show absorption at 280 nm for lignan compounds. Names of the proteins tested are shown on the left; corresponding protein IDs are listed in *SI Appendix*, Data S2. Peaks: a, (+)-pinoresinol (substrate of the reaction); b, (+)-piperitol; c, (+)-sesamin.

Analysis of publicly available RNA-seq data showed that the two *CcamCYP81Q* genes are expressed in seedlings and flower bud clusters (*SI Appendix*, Fig. S6) (27). Similarly, *CausCYP81Q111* and *CchiCYP81Q-1* were expressed in buds, ovaries, and seeds, but not in germinating seedlings (*SI Appendix*, Fig. S6) (19). These expression patterns are consistent with the observed accumulation of sesamin in the flower buds of *C. campestris* (Fig. 1). In contrast, *CcamCYP81AX6* showed negligible expression in these tissues.

Collectively, these results suggest that *Cuscuta* species synthesize sesamin through the enzymatic activity of PSSs encoded by *CYP81Q* genes, which are expressed predominantly in reproductive organs.

### Enlarged introns colonized by transposons in CYP81Q and other HGT genes

*SiCYP81Q1* has two exons separated by a single intron (12). We found that *Cuscuta CYP81Q* genes contain both more numerous and larger introns than those typically found in Lamiales, which commonly possesses two exons separated by a single small intron (Fig. 5A). The position of the second intron in *Cuscuta CYP81Q* genes is identical to that of the intron in *SiCYP81Q1* (Fig. 5A). These conserved intron positions between phylogenetically distant plants, i.e., *Cuscuta* and *Sesamum,* suggest a shared origin of these genes. Interestingly, transposons and their footprints were detected within the enlarged introns, including frequent insertions of a nonautonomous DNA transposon of the *Mariner*/*Tc1* class, *Stowaway* (*Sto*), and its associated footprints (Fig. 5A*, SI Appendix*, Data S7, Fig. S7). The highest sequence similarity was observed with *Sto* elements from *Ipomoea*, indicating that these elements originated from endogenous *Cuscuta* transposons rather than through horizontal transposon transfer (HTT) from other host plants. This supports the hypothesis that intron gain occurred after the HGT event in dodders.

**Figure 5.**
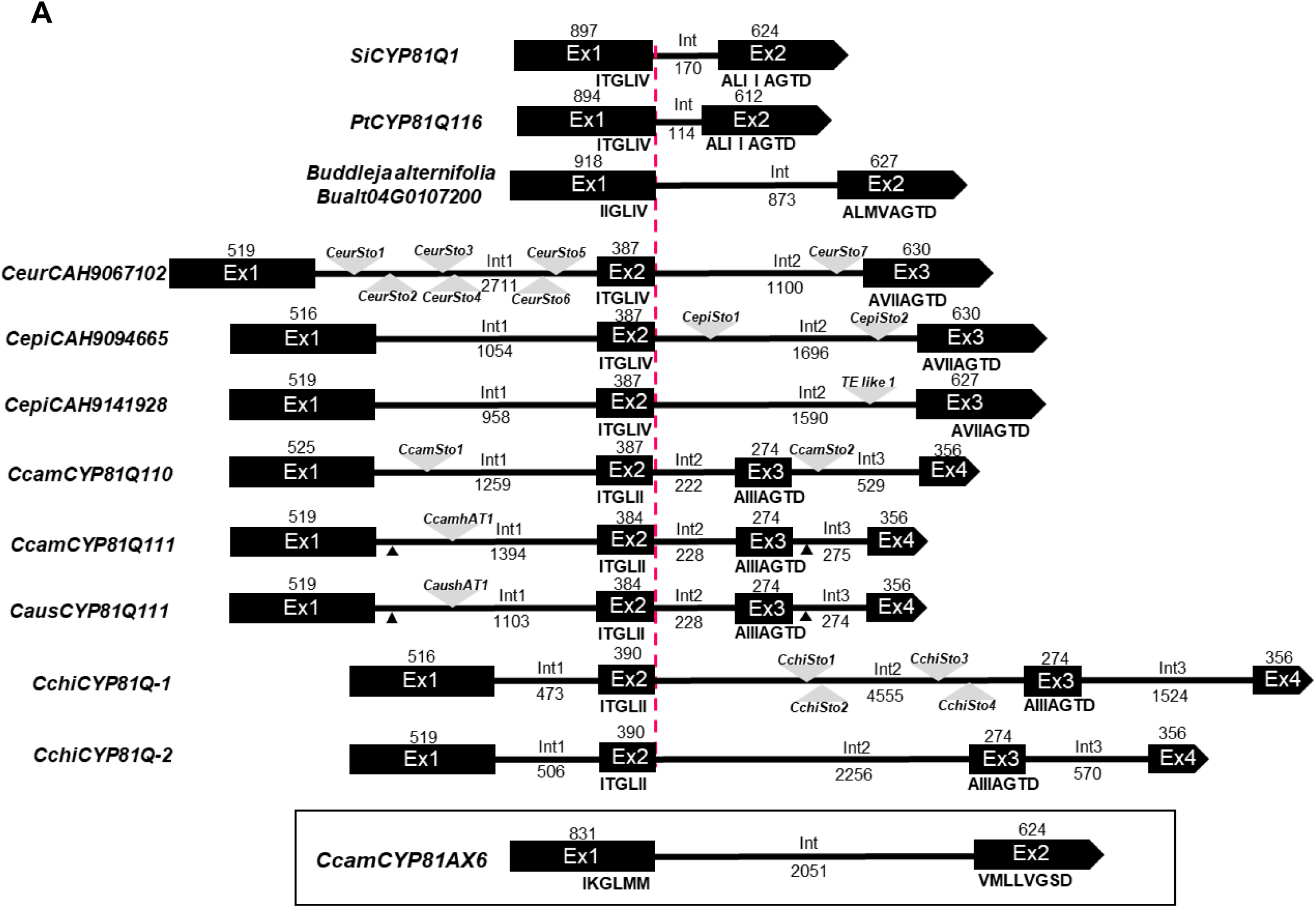
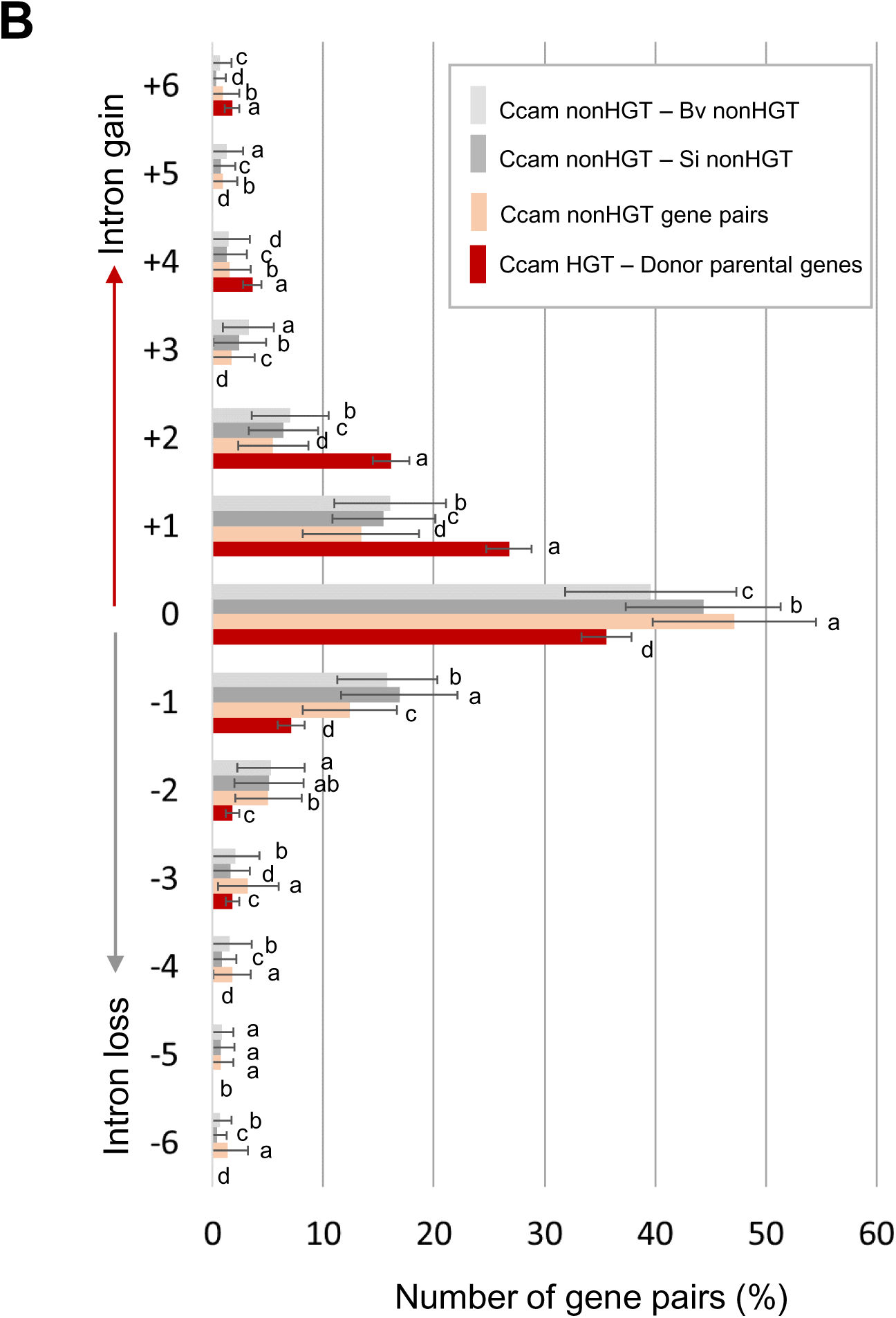
Genomic structures of CYP81Q genes: (A) Exon-intron structures of *CYP81Q* genes from Lamiales species (*Buddleja_alternifolia*_*BUALT_Bualt04G0107200*, *Handroanthus_impetiginosus_CDL12_01138*, *Paulownia tomentosa PtCYP81Q116*, *Phryma leptostachya PlCYP81Q38*, and *Sesamum indicum SiCYP81Q1*) aligned at the exon 1-intron (Ex1-Int) junction, and those from *Cuscuta* species aligned at the exon 2-intron (Ex2-Int2) junction. Amino acid sequences at the 3′-end of exon 1 or exon 2, and the 5′-end of exon 2 or exon 3 are shown. Ex: exon, Int: intron. Numbers indicate nucleotide lengths (bp). Grey triangles indicate transposons. Small black triangles indicate transposon remnants. *CcamCYP81AX6* has an exon-intron structure distinct from that of *CYP81Q* homologs. (B) Distribution of differences in intron numbers per gene. The vertical axis shows the difference in intron number between: Ccam non-horizontal transfer (nonHGT) genes and *Beta vulgaris* nonHGT genes with highest sequence similarity (nonHGT – Bv nonHGT); *Ccam* nonHGT genes and *S. indicum* nonHGT genes with highest similarity (Ccam nonHGT – Si nonHGT); *Ccam* nonHGT genes and their closest homologs (Ccam nonHGT gene pair); *Ccam* HGT genes and their orthologous genes from putative host species (Ccam HGT – Donor parental gene). The horizontal axis indicates the percentage of genes showing each level of intron number difference. Alphabetic letters above bars indicate statistically significant differences estimated by Tukey’s honest significant difference test (*p* < 0.05). (B) Distribution of differences in intron numbers per gene. The vertical axis shows the difference in intron number between: Ccam non-horizontal transfer (nonHGT) genes and *Beta vulgaris* nonHGT genes with highest sequence similarity (nonHGT – Bv nonHGT); *Ccam* nonHGT genes and *S. indicum* nonHGT genes with highest similarity (Ccam nonHGT – Si nonHGT); *Ccam* nonHGT genes and their closest homologs (Ccam nonHGT gene pair); *Ccam* HGT genes and their orthologous genes from putative host species (Ccam HGT – Donor parental gene). The horizontal axis indicates the percentage of genes showing each level of intron number difference. Alphabetic letters above bars indicate statistically significant differences estimated by Tukey’s honest significant difference test (*p* < 0.05).

Importantly, we observed that the position of *Sto* within the *CYP81Q* introns of *C. europaea, C. epithymum, C. campestris, C. australis,* and *C. chinensis* varies among species. On the other hand, the positions of the *hAT1* insertion and the *Sto* footprint within the *CYP81Q111* introns of *C. australis* and *C. campestris* exhibited striking similarity.

In response to the observed intron gain in *Cuscuta CYP81Q* genes, we investigated intron numbers in other 56 additional putative HGT genes in *C. campestris* (3). A high frequency of increased intron numbers (+1, +2, +4, +6) was observed in these HGT genes relative to their predicted donor orthologs (referred to as CcamHGT-donor parental genes; Fig. 5B*, SI Appendix*, Data S8). Conversely, no significant increase in intron number was observed between paralogous non-HGT genes in *C. campestris* (referred to as Ccam nonHGT gene pairs; Fig. 5B, *SI Appendix*, Data S8). To determine whether these intron gains reflect intrinsic interspecies differences, we compared intron numbers between *C. campestris* and predicted donor plants for orthologous gene pairs not derived from HGT. We examined 25 plant species identified as likely HGT donors (3), and present representative comparisons with *Beta vulgaris* and *S. indicum* (Ccam nonHGT-Bv nonHGT, Ccam nonHGT-Si nonHGT) (Fig. 5B*, SI Appendix*, Data S8). Neither comparison showed significant differences in intron numbers, indicating that the increased intron number in HGT genes is not attributable to baseline differences between donor and recipient plant species.

### Parasitism-evoked induction of SiCYP81Q1 in hosts and its transfer to parasites

Conserved genomic features and molecular functions of PSS genes in *Sesamum* and the distantly related parasitic *Cuscuta* plants suggest that HGT of *PSS* occurred through parasitism between their ancestors. To examine the feasibility of this model, we tested whether *C. campestris* is able to parasitize *S. indicum*. The results showed that *C. campestris* successfully established a parasitic connection to sesame (*SI Appendix*, Fig. S8 *A, B*), and developed flowers containing fertile seeds (*SI Appendix*, Fig. S8 *C, D*).

The presence of intron 2 in all of the *Cuscuta CYP81Q* genes (Fig. 5A) prompted us to investigate whether the putative HGT event involved the transfer of a genomic fragment containing an ancestral *CYP81Q* gene with an intron. To test this possibility, we analyzed sequence similarity between the genomic regions flanking *CYP81Q* genes in *Cuscuta* plants and *S. indicum*. While sequence similarity was observed between *CYP81Q* exon regions, the flanking regions showed extremely low similarity (*SI Appendix*, Fig. S9).

We then tested whether the acquisition of *CYP81Q* genes might have been mediated by RNA (28). Using *S. indicum* RNA-seq data (25, 26, 29), we conducted a search for intron-retained mRNA (IR-mRNA) of *SiCYP81Q1* and identified a minor IR-mRNA isoform (<3% of total transcripts) in *S. indicum* (*SI Appendix*, Fig. S10). Interestingly, the expression of both the spliced and IR forms of *SiCYP81Q1* was significantly induced by parasitization by *C. campestris* (*SI Appendix*, Fig. S8 *E-G*), and IR-mRNA of *SiCYP81Q1* was detected in the stem of *C. campestris* (*SI Appendix*, Fig. S8 *H*). These results suggest that a *CYP81Q* gene was horizontally transferred to *Cuscuta* via an HGT event associated with the induction of *CYP81Q* in the host plant in response to invasion by the parasitic plant.

## Discussion

### Convergent evolution of specialized lignans via HGT

Our data support an evolutionary scenario in which a functional *CYP81Q* gene was transferred from an ancestral plant in Lamiales, which is able to produce sesamin or related specialized metabolites, into an ancestral *Cuscuta* lineage through HGT. The conserved capacity of *CYP81Q* genes to form MDBs and generate specialized lignans suggests that this enzymic activity confers important physio-ecological advantages in dodders. At present, the frequency of functional parasitism-mediated HGT (pHGT) is difficult to estimate. Given that most horizontally transferred genes are likely to be eliminated from the parasite genome due to potential deleterious or non-beneficial effects on endogenous metabolism, the subset of conserved functional pHGT events may represent only the visible extent of a much larger number of unsuccessful pHGTs.

The estimated divergence time between Lamiales and Solanales is approximately 101.63 Ma (*SI Appendix*, Fig. S4), whereas the divergence between *Cuscuta* and *Ipomoea* species has been estimated around 50.38 Ma (*SI Appendix*, Data S6, Fig. S4). These estimates support the hypothesis that the *Cuscuta CYP81Q* gene was acquired via HGT after *Cuscuta* speciated from a common ancestor shared with *Ipomoea* (3). Interestingly, the monophyly of *Cuscuta CYP81Q* genes (*SI Appendix*, Fig. S3) suggests that the pHGT event occurred once in the most recent common ancestor (MRCA) of the subgenera *Cuscuta* and *Grammica*.

We identified *CYP81Q1* homologs in *C. campestris*, *C. australis*, *C. chinensis, C. epithymum*, *and C. europaea* (Fig. 2, *SI Appendix*, Data S1). These findings support the following conclusions: 1) the *CYP81Q* gene is widely conserved among species in the subgenera *Grammica* and *Cuscuta*; 2) the gene likely originated in a monophyletic ancestor located between the subgenera *Monogynella* and *Cuscuta*; and 3) the gene underwent changes in nucleotide sequence and modification of exon-intron structure from a common ancestral *CYP81Q*, in parallel with speciation events in *Cuscuta* and *Grammica*. Furthermore, the irregular distribution of *CYP81Q* genes in *Cuscuta* was observed in genomic regions having conserved gene synteny across *Cuscuta*, *Ipomoea*, and *Solanum* species (Fig. 3). These patterns support the hypothesis that *CYP81Q* genes were introduced into the *Cuscuta* genome via pHGT. Although we cannot fully exclude the possibility that an ancestral *I. nil* possessed a *CYP81Q1* ortholog that was subsequently lost, this scenario is considered unlikely as no members of the Convolvulaceae, other than *Cuscuta*, are known to accumulate sesamin or related specialized metabolites (7, 30).

Although numerous studies have suggested the possibility of HGT in various plant species (3,16–18), relatively little is known about the functional significance of HGT-derived genes in the recipient plants. In our study, *Cuscuta* CYP81Q proteins showed PSS enzymatic activity when expressed in yeast (Fig. 4). These findings demonstrate that the *Cuscuta CYP81Q* gene products are enzymatically active and catalyze a key branch-point reaction that converts pinoresinol, a common precursor of lignin and lignan biosynthesis, into sesamin and its downstream metabolic derivatives containing MDBs, such as water-soluble lignan cuscutoside (*SI Appendix*, Fig. S2).

We further evaluated the potential involvement of HGT in the evolution of specialized metabolism in dodder genomes by conducting Gene Ontology (GO) analysis. Notably, genes encoding pinoresinol/lariciresinol reductase (PLR), an enzyme that competes with PSS for the pinoresinol substrate and reduces it to secoisolariciresinol instead of oxidizing it form sesamin, were absent in dodder species but present in related taxa such as *Ipomoea* (*SI Appendix*, Fig. S11) (31). Conversely, PLR-mediated lignans such as arctigenin and matairesinol are present in *Ipomoea*, indicating the presence of functional PLR in these species (32). Given the reported competition between PLR and PSS for pinoresinol in engineered *Forsythia* species (33, 34), the loss of PLR in dodder may have facilitated lignan metabolic flux toward sesamin biosynthesis and its related metabolites following the HGT-mediated acquisition of PSS. These findings support the hypothesis that plant-to-plant HGT of *CYP81Q* genes represents a key evolutionary event, whereby a gene originally confined to Lamiales has become uniquely retained in *Cuscuta* within Solanales.

Nonetheless, the biological significance of sesamin and other specialized lignans in *Cuscuta* remains unclear. One possible function may be related to the durability of their seeds during dispersal (1, 2), potentially contributing to the widespread presence of dodder species. Whether HGT-mediated metabolic traits acquired from host plants has contributed to the unique physiology and ecology of *Cuscuta* remains an open question. Unraveling the genomic dynamics and biological functions of HGT genes in both parasitic dodders and their host plants could provide insights into how these parasites have adapted to persist in diverse environments on different continents.

### Origin of sesamin synthase genes

In contrast to *Cuscuta* species, we did not identify any homologs of *SiCYP81Q1* in the publicly available genomes of *Ginkgo biloba*, *Magnolia ashei*, or *Piper nigrum*, even though several phylogenetic relatives in Lamiales produce sesamin via *CYP81Q* genes (7, 8, 35) (*SI Appendix*, Fig. S1). This result is consistent with previous reports on cytochrome P450 phylogeny in plants, which indicate that *CYP81* family proteins are not conserved in basal angiosperms and gymnosperms (36). Given the substantial phylogenetic distance between these species and Lamiales, the sesamin synthase genes of these species, which are yet to be identified, are unlikely to share an origin with *SiCYP81Q1* (*SI Appendix*, Fig. S1). This likely contributes to the patchy phylogenetic distribution of sesamin production in the plant kingdom.

The high sequence similarity between *CcamCYP81Q111* and *CausCYP81Q111,* as well as their markedly lower similarity to *CcamCYP81Q110* (Fig. 2), motivated us to explore how *CYP81Q* genes diversified within the genus *Cuscuta*. The size of the *C. australis* genome (c.a. 264 Mbp) is approximately half that of *C. campestris* (c.a. 556 Mbp) (3, 19), suggesting that the *C. campestris* genome may have undergone doubling through either recent whole-genome duplication (WGD; autopolyploidization), estimated to have occurred around 1.5 Ma (3), or through divergence between the two subgenomes (BI and BII) of *C. campestris* around 3.13 Ma (*SI Appendix*, Fig. S4), whereas that between *C. australis* and *C. campestris* (B) occurred more recently, around 0.25 Ma, supporting a hybrid origin of *C. campestris* resulting from allopolyploidization between phylogenetically related lineages (3, 19, 31). This conclusion is consistent with previous taxonomic studies suggesting that *C. campestris* has a hybrid origin (2). Our findings on the systematic relationships of *CYP81Q* genes and *Cuscuta* genomes further support this hypothesis. *CcamCYP81Q111* and *CausCYP81Q111* share *hAT1* transposon in intron 1 and traces of *Sto* in intron 3, while *CcamCYP81Q110* contains *Sto* elements in introns 1 and 3 (Fig. 5A, *SI Appendix*, Data S7, Fig. S7). The genomic sequence flanking *CcamCYP81Q111* shows higher similarity to that flanking *CausCYP81Q111* than to that of *CcamCYP81Q110* (*SI Appendix,* Fig. S9). These results suggest that *CcamCYP81Q110* and *CcamCYP81Q111* are homeolog derived from a hybridization event between two *Cuscuta* species, each capable of sesamin biosynthesis, rather than resulting from WGD within an ancestral *C. campestris* genome. We additionally identified two *CYP81Q* genes from *C. epithymum* by genome mining, and from *C. chinensis* by genomic PCR. However, the markedly higher sequence similarity among these genes, along with the lack of genome assemblies for other species in the subgenus *Cuscuta*, precluded us from determining their possible hybrid origins (*SI Appendix,* Fig. S4). Future pan-genomic analyses of *Cuscuta* species will be necessary to clarify the evolutionary trajectories of this genus, which likely involve polyploidization, chromosomal rearrangement, and genome expansion (31, 37, 38). A chromosome-level assembly of the *C. campestris* genome has been recently reported (39), which will serve as a valuable reference for further refinement of the HGT analysis and *Cuscuta* phylogeny.

Our analysis of the sequence similarity between the genomic regions flanking *CYP81Q* genes in *Cuscuta* species and *S. indicum* revealed extremely low similarity (*SI Appendix,* Fig. S9). This observation implies that the pHGT of *CYP81Q* may have been mediated by host-to-parasite transfer of RNA. In Orobanchaceae, previously reported HGT events are hypothesized to involve the transfer of large genomic fragments, typically tens of kilobases in length (37). Similarly, the *C. campestris* genome contains conserved remnants of a horizontal DNA transfer from *Daucus carota*, spanning approximately 36 kbp (3). However, such a mechanism may not apply to *CYP81Q*. We experimentally confirmed that IR-mRNA of *SiCYP81Q1* is upregulated following parasitism by *C. campestris*, and that this transcript is translocated from *S. indicum* to *C. campestris* (*SI Appendix,* Fig. S8 *E-H*). These findings suggest the possibility that pHGT of *CYP81Q* may have occurred via the transfer of locally abundant mRNAs from host to parasite.

Although a previous study (16) showed that most pHGT events occur via DNA-based mechanisms, as evidenced by the presence of introns in most HGT genes, our findings on the transfer of IR-mRNA and the intron gain after HGT may obscure the original mode of transfer. The mechanisms underlying HGT remain unresolved, but those of HTT, involving both DNA and RNA transposons, may offer insights into the internalization of *trans*-genes (40).

### Transposon colonization in introns of HGT genes

The coding sequences of the *Cuscuta CYP81Q* gene have gradually diverged (*SI Appendix*, Data S9), and their genomic structures have undergone lineage-specific rearrangements, broadly reflecting the evolutionary time elapsed since the HGT event. Most importantly, *Cuscuta CYP81Q* genes acquired additional introns relative to the singlet intron present in the ancestral Lamiales *CYP81Q* genes (Fig. 5A). We compared the numbers of introns in 56 predicted HGT genes of *C. campestris* (from a previously reported set of 73 genes (3)) to those of their respective orthologs in putative host plants (3). The comparison revealed a consistent increase in the number of introns in HGT genes in *C. campestris* (Fig. 5B). This pattern is analogous to the substantial intron gain observed after the horizontal transfer of a bacterial amylase gene to Basidiomycetes (41), and represents the first report of extensive intron gain in HGT genes between parasitic and host plants. Among these HGT genes, intron gain occurred within protein-coding exons in nine genes, including *CYP81Q*. Conversely, 39 other genes exhibited intron gain within untranslated exons. These findings suggest that introns newly gained in protein-coding exons may have been retained under selective pressure in *Cuscuta*, because exon disruption could adversely affect protein function. On the other hand, the newly gained introns in untranslated exons may be retained under neutral selection, although such changes could still affect the efficiency of transcription or translation. Interestingly, the expanded introns in *CYP81Q* of *Cuscuta* species analyzed in this study were frequently colonized by transposons, particularly the nonautonomous DNA transposon *Sto* (42). The differential distribution of *Sto* elements among *CYP81Q* loci in various dodders suggests that transposon-mediated genomic rearrangements occurred repeatedly and independently in each lineage following the HGT event in the MRCA. This lineage-specific genomic restructuring of *CYP81Q* genes contrasts with the conserved intron structure observed in Lamiales, which retain a single short intron. Thus, understanding these processes is of particular interest in clarifying the mechanisms underlying the transposon colonization and intron gains following HGT.

Considering previous reports that DNA transposons known as *Introners* can generate introns in diverse eukaryotes (43-46), *Cuscuta Sto* elements may play a similar role in generating introns in pHGT genes. However, unlike *Introners*, the internal splicing motifs of *Sto* have not been shown to facilitate splicing (43, 44). The biological significance of intron gain is still a topic of debate. On one hand, the acquisition of introns can be deleterious, as it may increase the potential for splicing errors and associated mutations (47). On the other hand, intron gain may facilitate the integration of horizontally transferred genes by enabling their adaptation to the spliceosome of organisms with intron-rich genomes (41). Additionally, enlarged introns may provide a genomic context for the evolution of new functional elements, such as intronic non-coding RNAs (48). At present, it remains unclear whether intron gain in *Cuscuta* HGT genes confers harmful or adaptive effects. Therefore, clarifying the evolutionary significance of preferential intron gain by transposon colonization in these genes is an important subject for future studies. Beyond serving as an example of convergent metabolic evolution enabled by the acquisition of foreign genes, pHGT may also provide insights into the genomic mechanisms that govern the internalization and stabilization of foreign *trans*-genes in parasitic plant genomes.

## Materials and Methods

### Plant materials

*Cuscuta campestris* seeds were germinated and seedlings were parasitized onto host plants as described previously (49). *Nicotiana tabacum* plants were grown and parasitized by *C. campestris* as described previously (49). The *C. campestris*-*N. tabacum* parasitic complexes were grown at 25°C under a 16/8 h light/dark cycle, and flowers and fruits of *C. campestris* were harvested. The harvested tissues were frozen in liquid nitrogen and stored at -80°C. Seeds of *Sesamum indicum* cv. ‘Masekin’ were germinated and grown in potting soil (Sukoyakabaido, Yanmar, Osaka, Japan) mixed with the same volume of vermiculite (GS30L, Nittai Co., Ltd., Aichi, Japan) under natural sunlight from May to August in 2019 in the laboratory (Osaka Prefecture University, Japan).

Mature stems of 4- to 6-week-old *S. indicum* were parasitized by *C. campestris*. Stems of *S. indicum* and *C. campestris*, the parasitic interface, and *S. indicum* leaves were harvested at 6 days following parasitization, frozen in liquid nitrogen, and stored at -80°C until analysis.

Mechanical wounding of the *S. indicum* stem was performed with a razor blade and stems within 1.5 cm of the incision were harvested. Stems of *C. campestris*, measuring 10.5–13.5 cm in length and located at the parasitic interface, were harvested, washed twice with 70% ethanol for 2 min, and rinsed with nuclease-free water for 2 min to clean the tissue surface. *Cuscuta epithymum* and *C. europaea* were provided by Dr. J. Macas (Biology Centre, Czech Academy of Sciences). Specimens of naturally occurring *C. chinensis* and *A. indica* were harvested along the coast of Tokushima Prefecture, and along the shores of Lake Biwa in Shiga Prefecture, Japan, respectively.

### Chemicals

(+)-Sesamin (ChromaDex, Los Angeles, CA), (+)-sesaminol (Nagara Science Co., Ltd., Gifu, Japan), (+)-pinoresinol (Sigma-Aldrich Co. LLC, St. Louis, MO), and (+)-piperitol, synthesized previously (11), were prepared for use as the reference samples.

### LC-MS analysis

Lignans in extracts of flowers and seeds of *Cuscuta* plants (*C. campestris*, *C. europaea*, *C. epithymum*, *C. chinensis*, and *C. japonic*a) were analyzed as described previously (12, 26). A 10 mg sample of lyophilized flowers or seeds was homogenized to a fine powder using a TissueLyser II (Qiagen, Hilden, Germany). Next, 1 ml of 70% acetonitrile aqueous solution was added to the homogenized sample, followed by ultrasonic extraction at room temperature for 2 min. The resulting extracts were filtered and analyzed using an ion-trap time-of-flight mass spectrometer (LCMS-IT-TOF, Shimadzu Corp., Kyoto, Japan) equipped with a photodiode array detector (Shimadzu Corp.). Chromatographic separation was performed using a C18 column (TA12S03-1503WT, YMC Co., Ltd., Kyoto, Japan); 150 mm × 3 mm I.D., 3 µm particle size). The mobile phases were: A, 0.1% HCO_2_H-H_2_O; and B, 0.1% HCO_2_H-MeOH. A linear gradient of B was applied as follows: 30–50–90–90–30–30% B at 0–10–25–32–32.01–40 min, at a flow rate of 0.3 ml/min.

### PCR and reverse transcription-PCR (RT-PCR)

Genomic DNA from *C. campestris*, *C. japonica*, and *S. indicum* was extracted from stem tissue using a DNeasy Plant Mini kit (Qiagen), according to the manufacturer’s instructions. Total RNA was isolated using an RNeasy Plant Mini kit (Qiagen), according to the manufacturer’s instructions. The RNA samples were treated with either a DNase Set (Qiagen) or a TURBO DNA-free Kit (Thermo Fisher Scientific, Waltham, MA). After DNase treatment, cDNA was synthesized using an oligo dT primer and a PrimeScript RT Reagent Kit (Takara Bio Inc., Kusatsu, Japan), according to the manufacturer’s instructions. PCR products were amplified using gene-specific primer sets (*SI Appendix,* Data S10) as described previously (12). Briefly, genomic PCR, qPCR, and RT-PCR were conducted using PrimeSTAR Max DNA Polymerase (Takara Bio Inc.), GoTaq qPCR Master Mix, (Promega Corporation, Madison, WI) and PrimeSTAR GXL DNA Polymerase (Takara Bio Inc.), respectively.

### Genome and transcriptome

Publicly available *Cuscuta* transcriptome and genome datasets were used in this study. RNA-seq data for *C. campestris* were obtained from the DNA Data Bank of Japan Sequenced Read Archive (Accession number DRA009453; https://trace.ddbj.nig.ac.jp/dra/index_e.html/DRA009453) (27). The assembled genome sequence and annotations for *C. campestris* were obtained from the plaBi database (http://plabipd.de/portal/cuscuta-campestris) and for *C. australis* from the National Center for Biotechnology Information (NCBI) Sequence Read Archive (https://www.ncbi.nlm.nih.gov/bioproject/PRJNA394036). *CcCPR1*, *CcCYP81Q110*, *CcCYP81Q111*, *CcCYP81AX6*, and *CaCYP81Q111* correspond to contigs Cc043955, Cc046292, Cc015414, Cc047366, and RAL50776 (C65N0022E0.1), respectively. DNA-seq data for *C. americana* was obtained from SRA experiment ERR3569498 of BioProject PRJEB34450, and data for *C. californica* were obtained from SRA experiment ERR3569499. Synteny analysis was performed using BLASTP to identify best-hit protein sequences (*SI Appendix*, Data S5).

### Estimation of divergence time in *Cuscuta* species

To estimate the divergence times within *Cuscuta*, we first generated a pseudo-partition of the subgenomes of *C. epithymum*. Protein sequences of *C. epithymum* were subjected to a self– BLASTP search, and the resulting homology information was used to identify syntenic contigs with MCScanX (50). Syntenic contigs identified using MCScanX were similarly classified into subgenomes A and A’. Using OrthoFinder (51), we identified 472 single-copy orthologous genes shared across the proteomes of *C. australis* (GCA_003260385.1)*, C. campestris* (BI and BII) (31), *C. chinensis* (HI and HII) (31), *C. epithymum* (A and A’) (GCA_945859915.1), *C. europaea* (GCA_945859875.1)*, Ipomoea triloba* (GCF_003576645.1), *I. nil* (Asagao_1.2 genome, http://viewer.shigen.info/asagao/), *S. lycopersicum* (GCF_000188115.5), *S. indicum* (GCA_000512975.1), and *A. thaliana* (Araport11_pep_20220914 from TAIR, https://www.arabidopsis.org/). The amino acid sequences of the single-copy genes were aligned using MAFFT (--maxiterate 1000 --localpair) (52), and the resulting alignments were then converted to nucleotide sequences using PAL2NAL version 14.1 (53). From these alignments, 6649 fourfold degenerate third-codon transversion (4DTv) were extracted for divergence time estimation using BEAST version 2.7.5 (54). BEAUti was configured with the following parameters: Substitution Rate = “Tree” (estimated); Gamma Category Count = 4; Shape = estimated; Substitution Model = GTR; Clock Model = Relaxed Clock Log Normal; MCMC Chain Length = 20,000,000. The calibration points were set as described below: The divergence between A. thaliana (Brassicales) and Solanales: 85.80–128.63 Mya (55); The divergence between Solanaceae and Convolvulaceae: 59.1–83.9 Mya (56); The divergence between *Cuscuta* and *Ipomoea*: 47.8–56.0 Mya (57).

### Molecular phylogenetic analysis

The deduced amino acid sequences of *Cuscuta CYP81-*related genes used in this study are listed in *SI Appendix*, Data S3, and Data S4. The phylogenetic tree shown in Fig. 2 was constructed using the maximum likelihood (ML) method implemented in *SeaView* with PhyML (ln(L)=-23653.0; 1868 sites; GTR model with four rate categories) (58). The phylogenetic tree shown in *SI Appendix*, Fig. S3 was constructed using the BioNJ method in *SeaView* with 373 aligned sites and 1000 bootstrap replications (58). The timetree shown in *SI Appendix*, Fig. S6 was inferred using the *RelTime* method (59,60), applied to a phylogenetic tree with branch lengths estimated using the ML method and the Tamura-Nei substitution model. All evolutionary analyses were conducted using the *MEGA X* software package (61). Synteny analysis was performed based on BLASTP search results (cutoff, 1.00e-30).

### Genome comparisons

Genomic sequences spanning 5 kbp regions in both the upstream and downstream directions of *CYP81Q*-related genes were obtained based on the nucleotide sequences shown in Nucleotide ID of *SI Appendix*, Data S8. Pairwise similarities between these sequences were analyzed using *promer* in the *MUMmer* -v3.1.0 package (62). Structural similarities were visualized with *dotplots* using *mummerplot* (*SI Appendix*, Fig. S9).

### Molecular cloning

The nucleotide sequences of coding regions of *Cuscuta CYP81-*related genes used in this study are listed in Supplementary Data S9. cDNAs of the *CYP81-*related genes were obtained by RT-PCR from total RNA using gene-specific primers listed in *SI Appendix*, Data S10. *CaCYP81Q111* was artificially synthetized without codon-optimization (Eurofins Genomics, Luxembourg City, Luxembourg), based on the sequence of RAL50776. All P450 genes were cloned into yeast expression vectors and heterologously co-expressed in yeast with *C. campestris* cytochrome P450 reductase, *CcCPR1* (Cc043955), as described previously (26).

### Biochemical analysis

Biochemical analysis of *Cuscuta* CYP81Q proteins was performed essentially as described previously (26). Briefly, yeast cells expressing *Cuscuta CYP* genes were pre-cultured overnight at 30°C with rotary shaking at 120 rpm in 3 ml of synthetic defined liquid medium supplemented with the appropriate amino acids for each expression vector. Stationary-phase cultures (50 μl) were transferred into 1 ml of fresh medium in 24-well plates, supplemented with 100 µM of lignan substrates. The cultures were incubated for a further 24 h at 30°C with rotary shaking at 120 rpm. For extraction, cells were harvested together with the medium and disrupted by sonication. The homogenate (50 µl) was mixed with 50 µl acetonitrile and centrifuged at 21,000×*g* for 10 min. The supernatant was collected, filtered through a Millex-LH syringe filter (Merck Millipore, Burlington, MA), and subjected to HPLC analysis. Briefly, the filtered reaction products were separated using a Cortecs UPLC C18+ column (part# 186007401, 2.7 µm, 3 mm × 75 mm, Waters Corporation, Milford, MA). Chromatographic separation was performed with mobile phases A (0.1% trifluoroacetic acid-H_2_O) and B (0.1% trifluoroacetic acid-acetonitrile), using a linear gradient of 30–80–80–30–30% B at 0–1.4–1.8–2.0–2.5 min, at a flow rate of 1.25 ml/min. Lignan products were detected using a photodiode array detector at 280 nm.

### Preparation and imaging of agarose-embedded sections

Parasitic interface tissues were fixed in 4% (w/v) paraformaldehyde in phosphate-buffered solution (Fujifilm Wako Pure Chemical Corporation, Osaka, Japan), embedded in 8% (w/v) agarose, and sectioned into 200-μm sections using a MicroSlicer^TM^ ZERO 1N (DOSAKA, Kyoto, Japan). Histochemical staining of sections was performed using a 0.5% (w/v) aqueous solution of Toluidine Blue O (1B-481, Waldeck GmbH & Co., Munster, Germany). Stained sections were observed using a BX53 Biological Microscope (Olympus, Tokyo, Japan).

### Analysis of the number of introns per gene

A list of HGT genes in *C. campestris* and their putative species were obtained from previous literature (3). The number of introns per gene was determined from gene feature files associated with GenBank accession numbers listed in *SI Appendix*, Data S8. Gene pairs encoding proteins with the highest similarity were identified using BLASTP. Differences in intron number per gene were assessed using a randomly sample of 50 gene pairs, and the average frequencies were calculated using 2000 iterations of random sampling.

## Data Availability

The sequencing data are available from the DNA Data Bank of Japan under accession numbers PRJDB13351 and PRJDB15151.

## Acknowledgements

The authors thank P. Neumann and J. Macas (Czech Academy of Sciences) for providing us with fresh *C. epithymum* and *C. europaea* samples, and K. Krofta (Czech Hop Research Institute) for generously allowing us to use his laboratory facilities to prepare samples for lignan analysis. Computations were performed in part on the NIG supercomputer at the ROIS National Institute of Genetics. This work was supported in part by Grants-in-Aid for Scientific Research (18H03950 and 19H00944, JSPS to K.A.) and a Grant-in-Aid for JSPS Fellows (19J14848, JSPS to KS).

