## Supplementary Figures S1-11 for "Parasitism-mediated horizontal transfer of a functional cytochrome P450 gene entails transposon colonization in newly gained introns"

**This PDF file includes:**

Figures S1 to S11
Legends for Datasets S1 to S10
SI Methods
SI References

**Other supporting materials for this manuscript include the following:**

Datasets S1 to S10

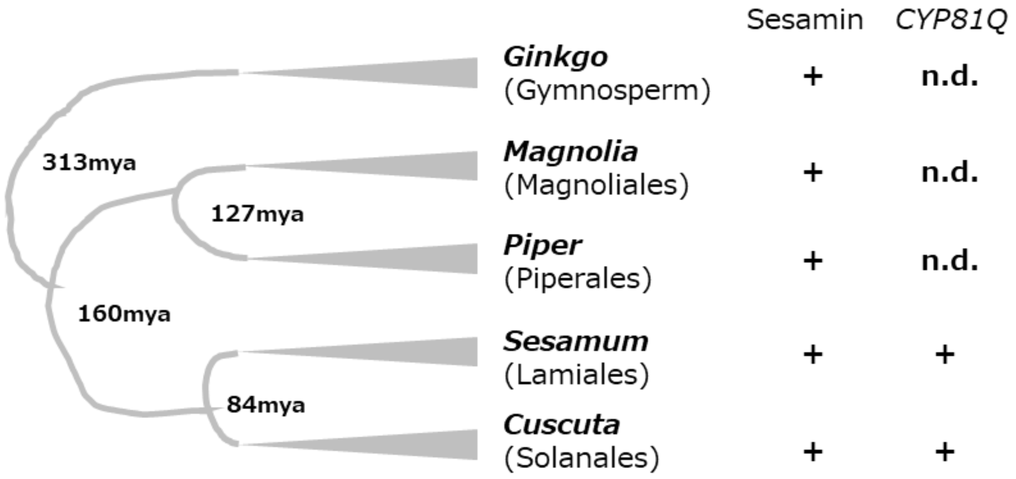

**Fig. S1.** Distribution of sesamin and *CYP81Q* gene in plant kingdom

Sesamin-producing plants are present sporadically in the plant phylogeny. Numbers on the tree indicate estimated divergent times according to the *Timetree* time scale of life28. "+" indicates detected. n.d. indicates not detected in a BLAST search querying *SiCYP81Q1* genes with default settings versus public genomes of *Ginkgo biloba* (GCA_024626585.1), *Magnolia ashei* (GCA_003571905.1), and *Piper nigrum* (Accession number PRJNA529758). MYA, million years ago.

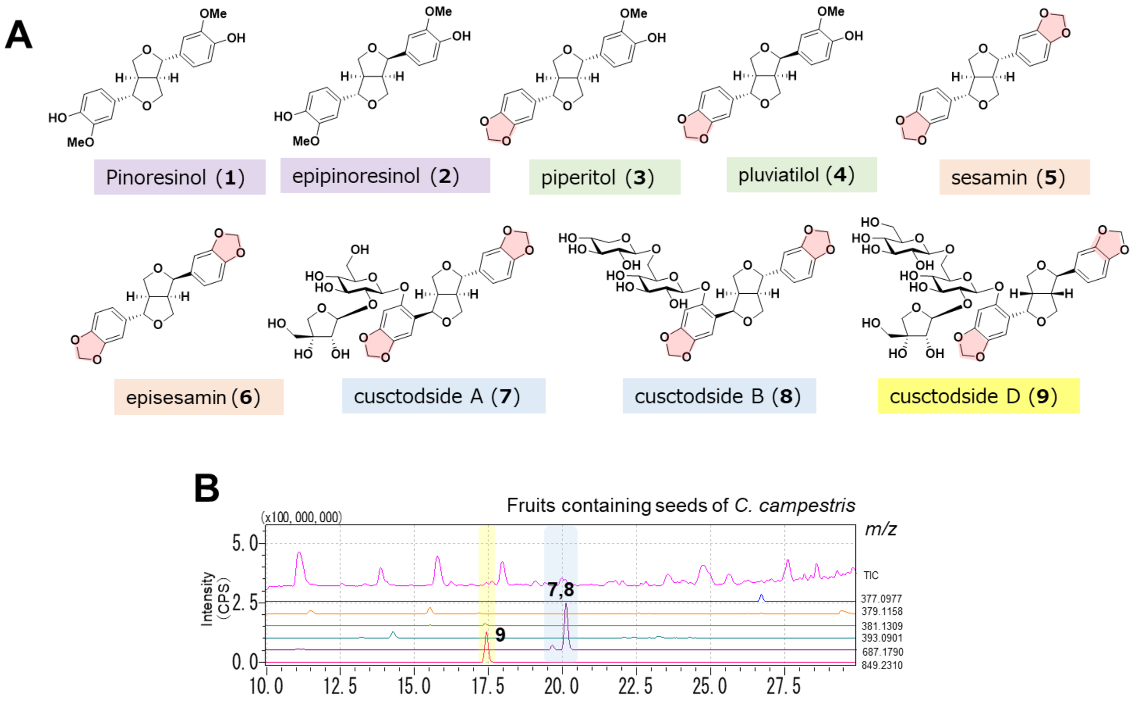

**Fig. S2.** Lignans in *Cuscuta* plants

**A.** Lignans detected in *Cuscuta* species. Lignan aglycons: (1) pinoresinol, (2) epipinoresinol, (3) piperitol, (4) pluviatilol, (5) sesamin, (6) episesamin, (7) cuscutoside A, (8) cuscutoside B, and (9) cuscutoside D. Lignans were detected using liquid chromatography-mass spectrometry (LC-MS) analysis. **B.** Cuscutoside A (7), cuscutoside B (8), and cuscutoside D (9), in addition to lignans detected in Figure 1, were detected in the seed-containing fruits of *C. campestris*.

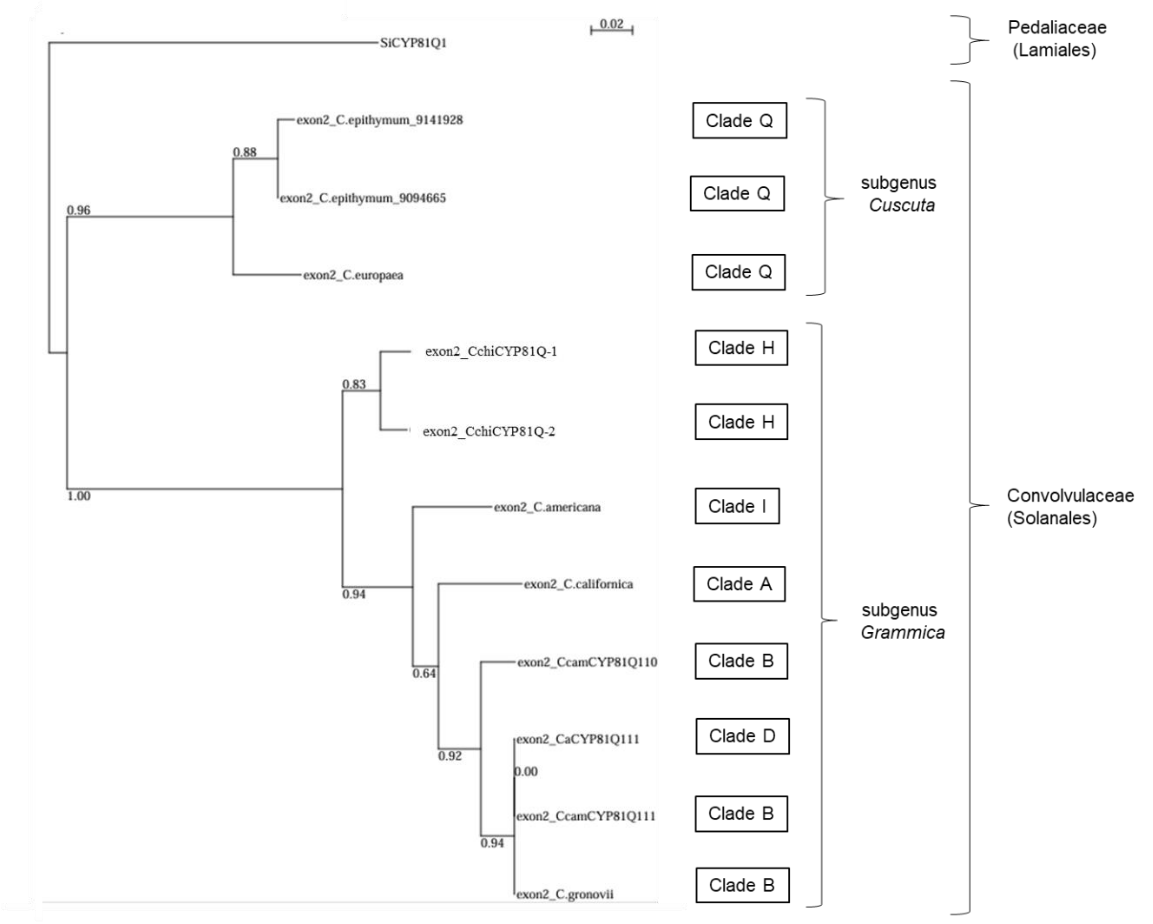

**Fig. S3.** Phylogeny of exon 2 of *CYP81Q* homologs

NGS data (SRA IDs ERR3569498, ERR3569499, SRR2143215, ERR3651373, and ERR3569501) for three *Cuscuta* spp. (*C. americana*, *C. californica*, and *C. gronovii*) were assembled using Trinity ver.2.9.1 (1). Contigs showing the highest similarity with *CcCYP81Q110* or *CcCYP81Q111* were selected by BLASTN search for each Trinity assembly. A phylogenetic tree was constructed by maximum likelihood using Seaview software (phyML: ln(L)=-1333.8, 387 sites, bootstrap=100 rep., GTR 4 rate classes) (2). The numbers at the branches indicate bootstrap values. The sequence of *SiCYP81Q1* exon 1, which corresponds to exon 2 of *Cuscuta CYP81Q*-related genes, was used as the outgroup. The exon 2 sequences from *C. americana*, *C. californica*, and *C. gronovii* that belongs to sugenus *Grammica* were co-clustered with those of *C. chinensis*, *C. australis* and *C. campestris*. The two sequences from subgenus *Cuscuta*, *C. europaea* and *C. epithymum*, formed a distinct clade from that of *Grammica*, roughly in line with the phylogeny of *Cuscuta* spp (3). The exon 2 nucleotide sequences for *CYP81Q*-related genes are available in *Supporting Information* Data S1. Phylogenetic clade (B to Q) is labeled with species names as defined in a previous phylogenetic study (3).

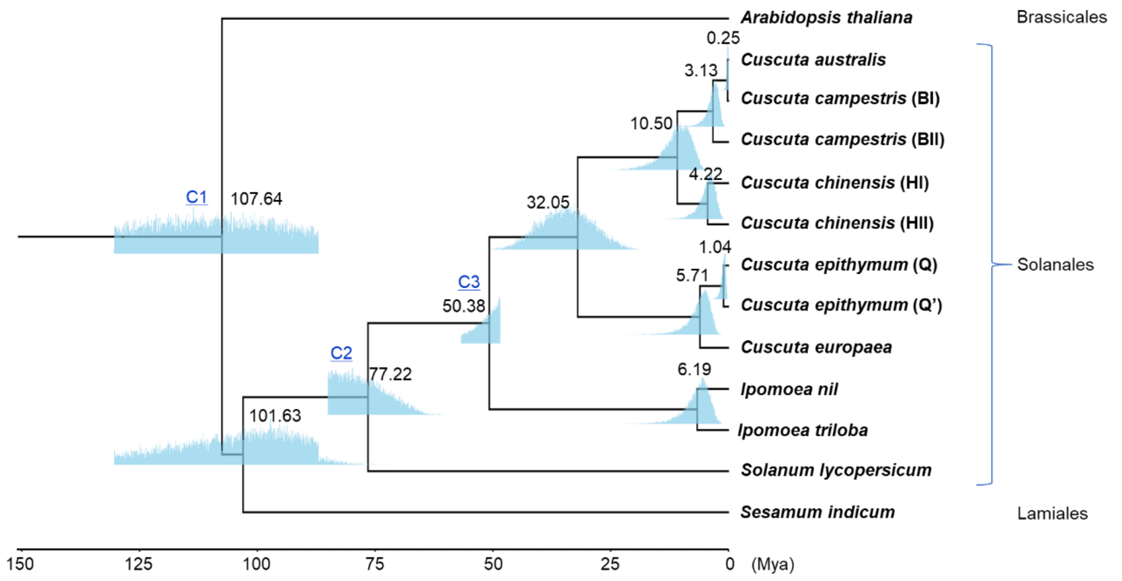

**Fig. S4.** Estimated divergent time of *Cuscuta* species

The blue shaded regions indicate the posterior distributions of node ages inferred from a Markov chain of 20,000,000 iterations. The first 20% of the samples were discarded as burn-in, resulting in 16,000,000 effective samples used for estimation. Median values are shown at each node. C1, C2 and C3: Calibration points.

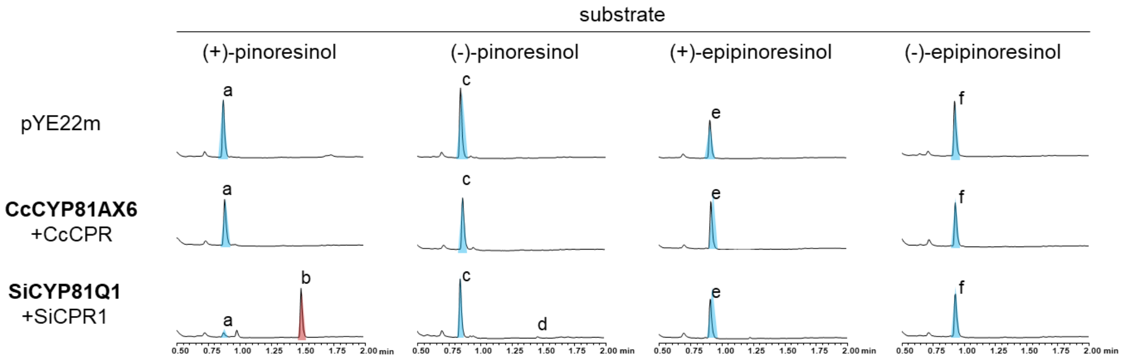

**Fig. S5.** Enzymatic activity of CcCYP81AX6

(Top) Negative control (empty yeast expression vector, pYE22m) for the bioconversion assay. Only the tested substrate (blue) was observed. (Middle) No product was observed in assays in which CcCYP81AX6 was co-expressed with CcCPR1. (Bottom) In reference assays in which SiCYP81Q1 was co-expressed with SiCPR1, (+)-pinoresinol was converted to (+)-sesamin (red), as shown in a previous report[14]. These results indicate that CcCYP81AX6 has no PSS activity. a, (+)-pinoresinol; b, (+)-sesamin; c, (-)-pinoresinol; d, (-)-sesamin; e, (+)-epipinoresinol; f, (-)-epipinoresinol.

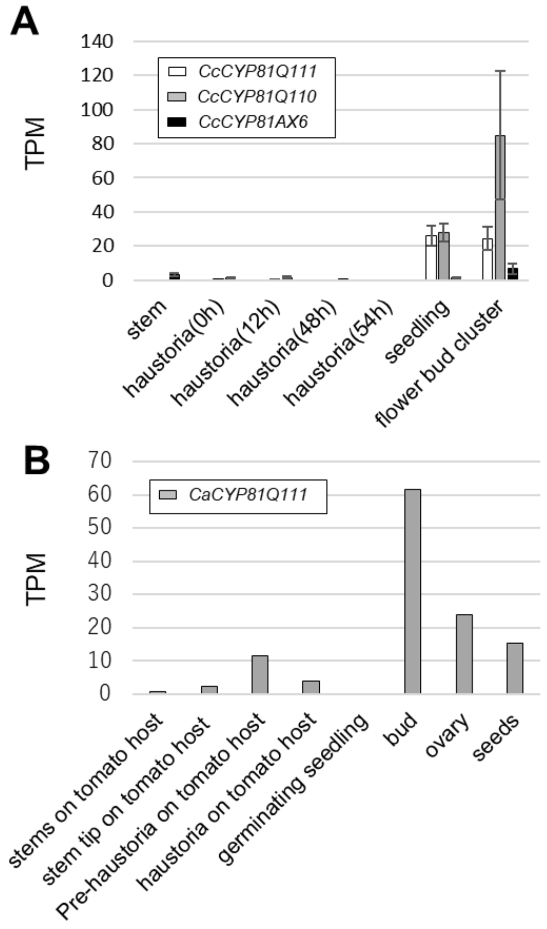

**Fig. S6.** Expression of *Cuscuta CYP81Q* genes in various tissues

**A.** RNA-seq analyses of *CYP81Q*-related genes in *Cuscuta campestris*. Each bar indicates the mean and standard deviation for three replicates. TPM values were estimated based on mapping of RNA-seq reads in DRA009453 (4) onto cucam_0.32.annot.cds.fasta from the plaBi database (http://plabipd.de/portal/cuscuta-campestris) (5). **B.** RNA-seq analyses of *CYP81Q*-related genes in *Cuscuta australis*. Each bar indicates the estimated TPM from a single-replicate RNA-seq experiment in PRJNA394036 (6). TPM, transcripts per million.

**A**

```
Intron 1
Cc_CYP81Q110    17   GTACTTT--GTA Cc_Sto1 TATTTCTTTT 284
Cc_CYP81Q111    17   GTACTATACATA TG      TATTTCTTTT  40
Ca_CYP81Q111    17   GTACTATACATA TG      TATTTCTTTT  40

Intron 3
Cc_CYP81Q110    119  TTGATATATA Cc_Sto2 TAGTCCTTTT   406
Cc_CYP81Q111    114  TTGACATATA CGGAG   TAGTGTTTTT   138
Ca_CYP81Q111    114  TTGACATATA CGGAG   TAGTGTTTTT   138
```

**B**

```
InSto1 5′ TIR  CTCCCTCAGTC
InSto1 3′ TIR  CTCCATCCGTC
Cc_Sto1 5′ TIR CTCCCTCCGTT
Cc_Sto1 3′ TIR CTCCATCCGTT
Cc_Sto2 5′ TIR CTCCCTCTGTC
Cc_Sto2 3′ TIR CTCCCTCCGTC
```

**C**

```
Cc_Sto1    CTCCCTCCGTTCTCAAAAATACTTCTTCTTTCATTT---------CCAACAAAACACTTC  51
Cc_Sto2    CTCCCTCTGTCCCAGAAAATCTTTCTCCTTTCTTTTTTCATCTGTCCCAAAAAACATTTC  60
           ******* ** *   *****  **** ***** ***         ** * ****** ***

Cc_Sto1    CACTTTCTATAAAAAA---ATACTTTACCCTTACTTTTTCATACCCTATCTATCACATCA 108
Cc_Sto2    TCATTTCTATTAAAAAGTCACTTTTACCCCTTACTTTTTCATACTCTACCTATCGCATCA 120
              ******* *****   *   **  ***************** *** ***** *****

Cc_Sto1    TATTTTTTCTCTAATAATC----TATCTATCACATCTTACTTTTACCCATCATATCAAGG 164
Cc_Sto2    TACTTTTTTACTAATAATCGATCTATCCATCAAATCATACTTTTACCCATCACATCAAAG 180
           ** *****  *********    **** **** *** *************** ***** *

Cc_Sto1    -ACAAAATTGGAAGCCAACTACAATTTAACACTTCCTCATTTTTACACTTGG---TCAAA 220
Cc_Sto2    GACATTTTTGTAAATTCAATACTAAAATACACTTCCCTTAAATCCCCCTTAAAAAACAAA 240
            ***   *** **    * *** *    ********      *  * ***      ****

Cc_Sto1    ATAAGAACTTTTTTTAGAACGGATGGAG                                 248
Cc_Sto2    ACTTTCATGTTTTCCGGGACGGAGGGAG                                 268
           *     *  ****   * ***** ****
```

**Fig. S7.** DNA transposons in introns of *Cuscuta CYP81Q* genes

**A.** Comparison of the intron 1 and 3 sequences. The TA dinucleotide target site duplication is indicated in boldface. The underlined sequences are the putative footprints of the *Stowaway* elements. Numerals indicate sequence positions in the intron sequences. **B.** Comparison of the conserved TIR sequences of *Stowaway* elements. **C.** Comparison of entire sequences of *CcSto1* and *CcSto2*. Terminal inverted repeat sequences are underlined.

**D**

Intron 1

| | | | | | | |
|---|---|---|---|---|---|---|
| *Cc_CYP81Q110* | 522 | AATTGATTGCCACTATGC | | | TTATTTTTTT | 549 |
| *Cc_CYP81Q111* | 270 | AATTGATTGT**AAGTGAGT** | *Cc_hAT1* | | **TAGTCAGT**TTAATTTTTT | 687 |
| *Ca_CYP81Q111* | 270 | AATTGATTGT**AAGTGAGT** | *Ca_hAT1* | | **TAGTCAGT**TTAATTTTTT | 719 |

**E**

| | | |
|---|---|---|
| *Tip100* 5′ | TIR | CAGGGGCGGAGGCA |
| *Tip100* 3′ | TIR | CAGGGGCGGAGCCA |
| *Cc_hAT1* 5′ | TIR | CAGTGGCGGAACCA |
| *Cc_hAT1* 3′ | TIR | CAGTGGCGGAGCCA |
| *Ca_hAT1* 5′ | TIR | CAGTGGCGGAACCA |
| *Ca_hAT1* 3′ | TIR | CAGTGGCGGAGCCA |

**F**

```
Cc_hAT1    CAGTGGCGGAACCACATGGGAGCAAATGACTACAATTGCAGGTAGTCCCTTTTCTATATA    60
Ca_hAT1    CAGTGGCGGAACCACATGGGAGCAAATGACTACAATTGCAGGTAGTCCCTTTTCTATATA    60
           ************************************************************

Cc_hAT1    TATATATATATATATATAAATAT-TATATATATATATATAT-----------------TA   102
Ca_hAT1    TATATATATATATATATATATATATATATATATATATATATATATATATATATATATATA   120
           ****************** **** *****************                 **

Cc_hAT1    TATATTATATTATAATATATATATATA-------------------ATATATATAGTACA   143
Ca_hAT1    TATATTATATATATATATATATATATATATATATATATATATATATATATATATAGTACA   180
           **********    *************                   **************

Cc_hAT1    AAGAGTACAAGCAAACCTAAAAAAAAGAGTACCGGGGAGGGAAAGAAGCACTACG---AC   200
Ca_hAT1    AAGAGTACAAGCAAACCTAAAAAAAAGAGTACCGGGGAGGGAAAGAG-CACTACGGACGC   239
           **********************************************  *******    *

Cc_hAT1    CGAGAGCAGAAAGAAGAATAGAAATACTCCTTATTTAAGATTACCATCCATGTCTACGAT   260
Ca_hAT1    GGAGAGCAGAAAGAAGAATAGAAATACTCCTTATTTAAGATTACCATCCATGTCTACGAT   299
           ************************************************************

Cc_hAT1    CAGAGTAAGGAGAAACCATTAGACCAATATTTATTTAAATATGTTCATCTCCGGTAACTG   320
Ca_hAT1    CAGAGTAAGGAGAAACCACTAGACCAATATTTATTTAAATATTTTCATCTCCGGTAACTG   359
           ****************** *********************** *****************

Cc_hAT1    CTAGATACGAACTTTTTCTTCTACTTTGCATCCACTGGTGTTGGATCTTGGCTCCGCCAC   380
Ca_hAT1    CTAGATACGAACTTTTTCTTCTACTTTGCAT-------TGTTGGATCTTGGCTCCGCCAC   412
           *******************************       **********************

Cc_hAT1    TG                                                             382
Ca_hAT1    TG                                                             414
           **
```

**Fig. S7.** DNA transposons in introns of *Cuscuta CYP81Q* genes (continued)

**D.** Comparison of the intron 1 sequences. Putative 8-bp target site duplication is indicated in boldface. Numerals indicate sequence positions in the intron sequences. Due to the low similarity of the target sequence region between *CcamCYP810Q110* and *CcamCYP810Q111* genes, it is possible that the *hAT* element was past inserted into the *CcamCYP810Q111* gene and exited. If this is the case, the underlined sequence could be a footprint sequence of the *hAT* element. **E.** Comparison of TIR sequences of *hAT* transposons from *Ipomoea purpurea* and *Cuscuta* plants. **F.** Comparison of entire sequences of the *hAT* elements from *Cuscuta* plants. Terminal inverted repeat sequences are underlined.

**G**

| | | |
|---|---|---|
| 141 GTAAATAC**TA** | *CeurSto1* (268 bp) | **TA**TCATTTTG 428 |
| 1021 TTGCAAAA**TA** | *CeurSto2* (193 bp) | |
| 1525 TTACCATA**TA** | *CeurSto3* (206 bp) | **TA** |
| | *CeurSto4* (246 bp) | **TA**TAACACTA 1998 |
| 2038 TATTTTAG**TA** | *CeurSto5* (207 bp) | **TA**ATATTTTT 2264 |
| 2269 ACCCCATA**TA** | *CeurSto6* (348 bp) | **TA**TATAATAA 2636 |

**H**

| | |
|---|---|
| *Stowaway* conserved TIR | CTCCCTCCGTT |
| *CeurSto1* 5′ TIR | CTCCCTCCGTC |
| *CeurSto1* 3′ TIR | CTCCCTCCATC |
| *CeurSto2* 5′ TIR | CTCCCTCCCTT |
| *CeurSto2* 3′ TIR | |
| *CeurSto3* 5′ TIR | CTCCCTCCGTC |
| *CeurSto3* 3′ TIR | CTACCTCCGTC |
| *CeurSto4* 5′ TIR | CTCCCTCCGTC |
| *CeurSto4* 3′ TIR | CTCCCTCCGTC |
| *CeurSto5* 5′ TIR | CTCCCTCCGTT |
| *CeurSto5* 3′ TIR | CTTCCTCCGTT |
| *CeurSto6* 5′ TIR | CTCTCTCCGTT |
| *CeurSto6* 3′ TIR | CTCCATCCGTT |

**I**

```
CeurSto2   CTCCCTCCCTTCTTATTTTATCTTCCAAATTGGTTTGTCACACAATTTAAGATAGTGAAT   60
CeurSto5   CTTCCTCCGTTCCTATTTTATCTTCCAAATAGGGTTGGCACACAATTTAAGATAGTGGGT   60
           ** ***** *** ***************** ** *** *******************  *

CeurSto2   ATTGTGTGGTGGAGAGTTGTGTAGAGGAAAGAGAAATGTGTAAGAAATGTTGTGAACCAC  120
CeurSto5   ATTGTGTGGTGGAGAGTTGTGCAGAGGAGAGAGAAATGTGTAAGAAATGTTGTGAACCAC  120
           ********************* ****** *******************************

CeurSto2   TAATTCTTTACTAAAAAGGGAAAGTGAAAGTTTTAATTAGAACAGACGAAAAAGGAAAAT  180
CeurSto5   AAATTCTTTACTAAAAAGGGAAAATGTAAGTTTTAATTGGAACGAACGAAAAAGGAAAGT  180
            ********************** ** *********** ****  ************* *
CeurSto2   TAGAAGTTTTAAT--------------                                   193
CeurSto5   TGGAAGTTTTAATAGGAACGGAGGGAG                                   207
           * ***********
```

**Fig. S7.** DNA transposons in introns of *Cuscuta CYP81Q* genes (continued)

**G.** The insertion sites of the *CeurSto* elements. The TA dinucleotide target site duplications are indicated in boldface. *CeurSto3* and *CeurSto4* are adjacent to each other across 2 bp of TA. *CeurSto2* lacks its 3’ terminus. Numerals indicate sequence positions in the intron sequences. **H.** Comparison of the conserved TIR sequences of *Stowaway* elements. **I.** Comparison of entire sequences of *CeurSto2* and *CeurSto5*. *CeurSto2* and *CeurSto5* are inserted in opposite directions, and the complementary sequence of *CeurSto5* is shown. The conserved sequences of the terminal inverted repeats are underlined.

**J**

```
CeurSto7              ---------CGTCCCGTTGGATTTTGTCAAGTGCAAAGTATGTGAAAAATAAGAATAGTA   51
3002310-3002510       TACTACCTCCGTCCCGGTGGACTTTGCCAAGTGCAAAGTAGGTGAGAAATAAGAATAGTA   60
25554319-25554519     TACTACCTCCGTCCCGGTGGACTTTGCCAAGTGCAAAGTAGGTGAGAAATAAGAATAGTA   60
37127923-37127724     TACTACCTCCGTCCCGGTGGATTTTGCCAAATGCAAAGTAGGTGAGAAATAAGAATAGTA   60
                               ******* **** **** *** ********* **** **************

CeurSto7              CGAAGTACAAAGTGTGTGATTAAGATTTGTGTAGAGATGAGAGAAATGTACACAACCATA   111
3002310-3002510       A-------AAACTGTGTGGTTGAGATTTGTGCAGAGGAGAGAGAAATGTGCAGAACCACA   113
25554319-25554519     A-------AAACTGTGTGATTGAGATTTGTGCAGAGGAGAGAGAAATGTGCAGAACCACA   113
37127923-37127724     A-------AAACTGTGTGGTTGAGATTTGTGCAGAGGAGGGAGAAATGTGCAGAACCACA   113
                              *** ****** ** ********* ****  * ********* ** ***** *

CeurSto7              AATTAGTTGCTTTAATAGGAAAGTGAGAAGTTTTTTGTGACAATCAAAAAAGAAAAGTT   171
3002310-3002510       AATTAGTTGTTTTAATAGGAAAGTGACAAAGTTTTTTGGGACAACCAAAAAAGGAAAGTT   173
25554319-25554519     AATTAGTTGCTTTAATAGAAAAGTGCCAAAGTTTTTTGGGACAACAAAAAAAGGGAAAGTT   173
37127923-37127724     AATTAGTTGCTTTCATAGGAAAGTGACAAAAATTTTTGAGACAAC-AAAAAAGGAAAGTT   172
                      ********* *** **** ******  **   ****** *****  ***** * ******

CeurSto7              GGCAAAGTCCAATGGGACG---------                                   190
3002310-3002510       GGCAAAGTCCAATGGGACGGAGGGAGTA                                   201
25554319-25554519     GGCAAAGTCCAATGGGACGGAAGGAGTA                                   201
37127923-37127724     GGCAATGTCCAATGGGACGGAGGAAGTA                                   200
                      ***** *************
```

**Fig. S7.** DNA transposons in introns of Cuscuta CYP81Q genes (continued)

**J.** Comparison of *CeurSto7* and its relative sequences on the *C. europaea* genome sequence (CAMAPE010000005.1). The titles of sequences other than *CeurSto7* are positions in the genomic sequence. *CeurSto7* lacks its TIR sequences. The conserved sequences of the terminal inverted repeats are underlined. The target site duplication sequences are shown in boldface.

**K**

| | | | | | |
|---|---|---|---|---|---|
| CAH9094665 | 180 | AGAAATTA**TA** | *CepiSto1* (253 bp) | **TA**TGTGGGTT | 452 |
| CAH9141928 | 180 | AGAAATTATA | | --TGTGGGTT | 197 |
| CAH9094665 | 1259 | TTCACA--------- | *CepiSto2* (247 bp) | TAGTTGTGTT | 1521 |
| CAH9141928 | 1386 | TTCACA(18 bp)TA | | --CTTGTGTT | 1419 |
| CAH9094665 | 1051 | GAAGTTGAATTATTAT |  | ---------TGAATGAT | 1074 |
| CAH9141928 | 815 | GAAGTTG**AATTATTAT** | *TE like1* (362 bp) | **AATTATTAT**TGAATGAT | 1209 |

**L**

| | |
|---|---|
| *Stowaway* conserved TIR | CTCCCTCCGTT |
| *CepiSto1* 5´ TIR | CTTCATCCGTT |
| *CepiSto1* 3´ TIR | CTCCCTCCGTT |
| *CepiSto2* 5´ TIR | ---------TT |
| *CepiSto2* 3´ TIR | CTTCCTCTGTT |

**Fig. S7.** DNA transposons in introns of *Cuscuta CYP81Q* genes (continued)

**K.** Comparison of the intron 2 sequences of *C. epithymum CYP81Q* genes. Shown in bold are target site duplications of 2 bp and 8 bp for *CepiSto1* and *TE like1*, respectively. Numerals indicate sequence positions in the intron sequences. *CepiSto2* is deleted at its 5´ terminus along with its adjacent sequence. **L.** Comparison of the TIR sequences of *CepiSto* elements.

**M**

```
CepiSto2        -----------TTCTTTTTTAATTGCAACATTGGACTTTTTGCACTACTCACATATTCTA    49
972282-972539   TACTACCTCCGTTCTCTTTTAATTGCAACATTGGACTTTTTGCACTATTCACATGTTCTA    60
166873-167132   TACTACCTCCGTTCTCTTTTAGTTGCAACGTTGGACTTTTTGCACTATTCACATATTCTA    60
1374411-1374670 TACTACCTCTGTTCTCTTTTAATTGCAACGTAGGACTTTTTGTACTATTCACACATTCTA    60
                           **** ***** ******* * ********** **** *****  *****

CepiSto2        CTTTGACTATATTTTATTTATTAATGTATTATAAATTAAATTTTTAAAAAATATAGTTAA   109
972282-972539   CTTTGACTATATTTTATTAATTAATGTTTTATAAATTAAAATGTTAAAAAATATTGTTAA   120
166873-167132   CTTTGACTACATTTTATTAATTGATGTATAATAAATTAAATTTTTAAAAAATATTGTTAA   120
1374411-1374670 CTTTGAGTATAGTTTATTTATTAATGTATTACGAATTTAATTTTTAAAAAATATTGTTAA   120
                ****** ** * ****** *** **** * *  **** ** * *********** *****

CepiSto2        ATTATTCACATTCTAAATTTTCGAAATCTGGATTTTTTCAATTTTTTACTAATAAGGAAC   169
972282-972539   ATTCTTCTCATTCTAAATTTTCAAAATATGAATTTTTAAATTTTTTTACTATTAAGGAAT   180
166873-167132   ATTCTTCTCATTCTAAATTTTCAAAATATGAATTTTTTATTTTTTTTACTAATACGGAAT   180
1374411-1374670 ATTCTTCTCATTCTAAATTTTCAAAATATGAATTTTTGAAAATATTTACTAATAAGGATT   180
                *** *** ************** **** ** ******     * ******* ** ***

CepiSto2        GAAAGATGTTAATGATCAAACTTGTGCGTTGGCAAACGTGAAATATTGACTGTTGCAATT   229
972282-972539   GAAAGATATTAATGGTCAAACTTGTGCGTTGGCAAACGT--AATGTTGACTGTTGCAATT   238
166873-167132   GAAAGATATTAATGGTCAAACTTGTGCGTTGGCAAACGTGAAATGTTGACTGTTGCAATA   240
1374411-1374670 GAAAGATATTAATGGTCAAACTCGTGCGTTGACAAACGTGAAATGTTGACTGTTGCAATT   240
                ******* ****** ******* ******** *******  *** **************

CepiSto2        AAAAGAGAACAGAGGAAGTA                                           249
972282-972539   AAAAGGGAACGGAGGAAGTA                                           258
166873-167132   AAAAAGGAACGGAGGAAGTA                                           260
1374411-1374670 AAAAGGGAACGGAGGAAGTA                                           260
                ****  **** *********
```

**N**

```
TE like1 5´ TIR      GGAAAAATTGATTATAATAATCCCAACTATTGACGGTCCGCTTATAATAATCTCAACTGT     60
TE like1 3´ TIR      GGAAAAATCGAAAATAATAATCCCAACTATTACCCATCCGCTAACATTAATCCCAACTAT     60
                     ******** **  ******************  *  ****** * * ***** ***** *

TE like1 5´ TIR      CGATTATTTCAGAATAATCCCAACTTTAGGGACCTTCTGCCAATAATAGTCC            112
TE like1 3´ TIR      TGATTATTTTTGTATAATCCCAACTATATGGGTCATCCGCCAATAATGGTCC            112
                      ********  * ************ ** **  * ** ********* ****
```

**Fig. S7.** DNA transposons in introns of *Cuscuta CYP81Q* genes (continued)

**M.** Comparison of *CepiSto2* and its relatives on the *C. epithymum* genome sequence (CAMAPF010000082.1). The sequence titles other than *CepiSto2* are positions in the genomic sequence. *CepiSto2* lacks its 5´ TIR. The conserved sequences of the terminal inverted repeats are underlined. The target site duplication sequences are shown in boldface. **N.** The TIR sequences of *TE like1*.

**O**

*Cchi_CYP81Q2* intron 2

| | | | | |
|---|---|---|---|---|
| 2,260 | AATTGTAA**TA** | *CchiSto1* | **TA**TTACTTTA | 2541 |
| 3,147 | GAATATAA**TA** | *CchiSto2* | **TA**TTAGATTA | 3382 |
| 3,682 | TAAGATAG**TT** | *CchiSto3* | | |

**P**

| | |
|---|---|
| *Stowaway* conserved TIR | CTCCCTCCGTT |
| *CchiSto1* 5′ TIR | CTCCCTCTGTC |
| *CchiSto1* 3′ TIR | CTCCCCCCGTC |
| *CchiSto2* 5′ TIR | TTACCTCCGTC |
| *CchiSto2* 3′ TIR | CTCCCTCCGTC |
| *CchiSto3* 5′ TIR | CTCCCTCCATC |

**Q**

```
CchiSto4   3825  GGAAAGTGGAACCTTTCTTTGCGACAAACAAAAAAGAAAA  3864
CceurSto7  129   GGAAAGTGAGAAGTTTTTTTGTGACAATCAAAAAAGAAAA   168
                 ********  *  *** **** ***** ************
```

**Fig. S7.** DNA transposons in introns of *Cuscuta CYP81Q* genes (continued)

**O.** The target sites of the *Stowaway* elements in intron 2 of *CchiCYP81Q-2*. The TA dinucleotide target site duplication is indicated in boldface. Numerals indicate sequence positions in the intron sequences. **P.** Comparison of the conserved TIR sequences of *Stowaway* elements. **Q.** Comparison of entire sequences of the partial *Stowaway* elements of *CchiSto4* and *CeurSto7*. Numerals indicate the position within the intron for *CchiSto4*, and for *CeurSto7*, it denotes its position within its own sequence.

**R**

*Cchi_CYP81Q2* 228 TATGGGTAAAGTAAT *CchiLTR1* ATGTTTTTTAAGCGG 953

*CcLTR1* scaffold24 875183 TTGTTTGACA**TATAC** *CcLTR1* **TATAC**TTTCTCTTGA 874458

**S**

```
CcLTR1      TGTAACACCCCGAC---CCCCACCCTAG-CCGTAATTGTCCGCTCATAGGATTGTAGATT    56
CchiLTR1    TGTAACACTCTAGCTAGCCCCACCCCAGGCCGGTAATGACTACTCATGAGATTATAGGTT    60
            ******** *   *   ******** ** ***  * ** *  *****  **** *** **

CcLTR1      GGCCCCACAAACCAACACAAGTCTTTTCCGCGCACTTTGTCCTCACTCGTGCGCGCCCAA   116
CchiLTR1    GGCCCTACAATTCAACACAAGATTTTCTCGTGCATTTTGGCCTCACTCATGCACGCTCT-   119
            ***** ****  *********  ***  ** *** **** ******** *** *** *

CcLTR1      GGAAACTTCCCAGGGGGTCACCCATCCTAGAATTGCTCCAAGTCAAACACGCTTAACTTT   176
CchiLTR1    GAAAACTTCCCAGGGAGTCGCA-ATCCTAGAATTGCTCCAAATTAAACATGCTTCACCTT   178
            * ************* *** *  ****************** * ***** **** ** **

CcLTR1      GGAGTTTTTAGGTGATGAGCTCTCGAAAAGAAGATGCACCTTGTTGGTAT--GAGTAGTA   234
CchiLTR1    GGAGTTTTTAAGTTATGAGCTCTTGAAAAGAAGGTGCATCTTGTTGATATACGAGTAGTA   238
            ********** ** ********* ********* **** ******* ***  ********

CcLTR1      CTATCAATCCATTTGAGCTCTTCTCATTC----TCATCTTAACACAGGGTATTACATTCA   290
CchiLTR1    TAATCAATTTTTTTGAGTTCTTTTCAATCCATCTCATTTCAACACGGGATATTACATTCA   298
              ******   ****** **** *** **    **** * ***** ** ***********

CcLTR1      CCCCCACTTAGGAATGCAACGTCCTCGTTGCATCCTTTTTTTAAACCCCTAGATCTTCCC   350
CchiLTR1    CCCTCACTTAAGAATACAACATCCTCGTTGCAC-----------GCCCCAAGATCCTCCC   347
            *** ****** **** **** ***********             **** ***** ****

CcLTR1      C-GCTAGGTGAACCAAGCTCTTCTTGGCCCGTCACGGTCGCAGGGTTGCTCTGATACCAT   409
CchiLTR1    CCACTAGGTGAACTGAGCTCGT--TGGCCCATCACAACCGCAAAATTGCTTTGATACCAT   405
            *  **********  ***** *  ****** ****   ****   ***** *********

CcLTR1      TTGTAACACCCAGGCCCCCACCCTAGCCGTAAGTGTCCGCTCATAGGATTGTAGACTGGC   469
CchiLTR1    TTGTAACACTTTGGCCACCCCTCGGGCCAGAAGTGACTACTCATGAGATTGTAAACTTGC   465
            *********   **** ** * *  ***  ***** *  *****  ******* *** **

CcLTR1      CCCACAAACCAACACAAGTCTTTTCCGCGCACTTTGTCCTCACTCGTGCGCGCCCAAAGA   529
CchiLTR1    CTCACGAACCAACACAAGTCTTTCCCGCGCATTTTAGCCTCATTCATGTGCGCCTTGA-A   524
            * *** ***************** ******* ***  ***** ** ** *****   * *

CcLTR1      AACTTCTCAGGGGGTCACCCATCCTAGAATTGCTCCAGGTCAAGCACGCTTAACTTTGGA   589
CchiLTR1    AACATTCCAGGGGGTCACCAATCATAGAATTGCTCCAAGTCAAGCATGTTTTACTTTGGA   584
            *** *  ************ *** ************* ******** * ** ********

CcLTR1      GTTTCTAGGTGATGAGCTCTCGAAAAGAAGATGCACCTTGTTGGTATGAGTAGTACCATC   649
CchiLTR1    GTTTCTAAGTTATGGACTCTCAAAAATTAGATGCATCTTGTTGGTAAAAGTGGTACTATA   644
            ******* ** ***  ***** ****  ******* **********  *** **** **

CcLTR1      AATCCATTTA--AGCTCTTCTCAT---TCTCATCTTAACACAGGGTATTACA           696
CchiLTR1    AATCCTTTTTTTAGTTCTTTCAATCCATCTCATTTCAATTCGGAGTATTACA           696
            ***** ***   ** ****   **   ****** * **  * * ********
```

**Fig. S7.** DNA transposons in introns of *Cuscuta CYP81Q* genes (continued)

**R.** The target sites of the retrotransposons, *CcamLTR1* and *CchiLTR1*. *CcamLTR1* is identified in the *C. campestris* genome assembly, Ccam0.32_scaffold24 (GenBank accession number: OOIL02001568.1). The putative 5-bp target site duplication is highlighted in boldface. Numerals denote sequence positions in the intron sequences and the scaffold sequence for *CcamLTR1* and *CchiLTR1*, respectively. **S.** A comparison of the entire sequences of the retrotransposons is shown, with the Long Terminal Repeat (LTR) sequences being underlined.

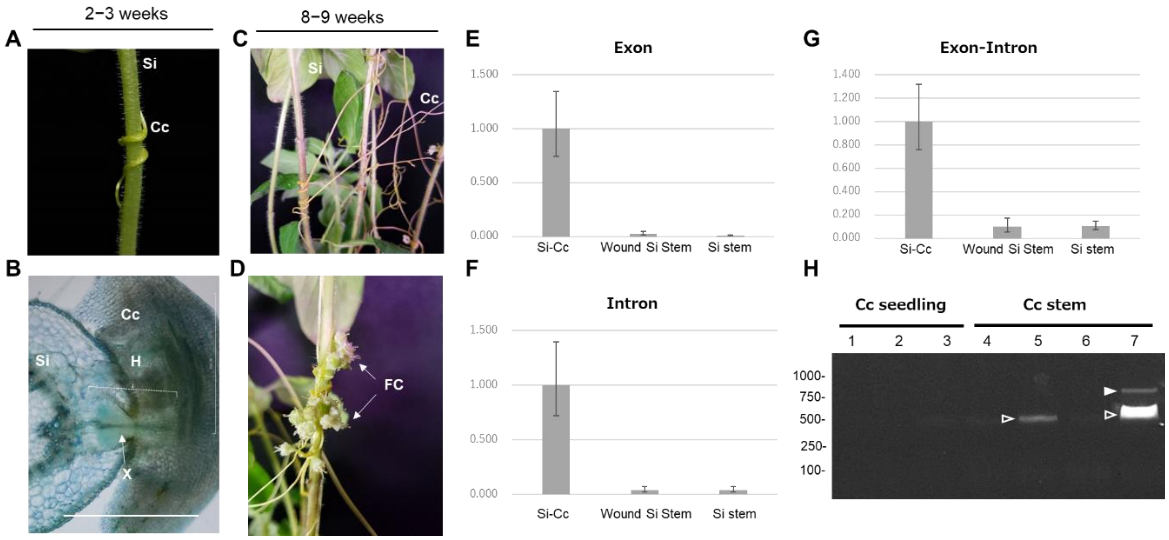

**Fig. S8.** Parasitism of *Cuscuta campestris* on *Sesamum indicum* and transfer of *SiCYP81Q1* mRNA

**A.** Parasitism of the parasite, *Cuscuta campestris* (Cc), on the host, *Sesamum indicum* cv. Masekin (Si), at 2-3 weeks after germination. **B.** Cross section (thickness=200 μm) of haustorium with toluidine blue staining at 2-3 weeks after germination. Cc, *C. campestris*; Si, *S. indicum*; H, haustorium; X, haustorial xylem. Scale bar, 500 μm. **C.** *C. campestris* (Cc) on *S. indicum* cv. Masekin (Si) at 8-9 weeks after germination. **D.** *C. campestris* formed a flower and developed fertile seeds. FC, floral cluster of the parasite. Gene expression of *Si_CYP81Q1* (**E.** exon, **F** intron, **G.** exon-intron) in stems of *S. indicum* at the parasitic interface of *C. campestris* (left), at mechanical wounded site (middle) and non-treatment (right). **H.** RT-PCR detection of *Si_CYP81Q1* in the non-parasitizing stems (Cc seedlings) and parasitizing stems of *C. campestris* (Cc stem). Each lane indicates an independent replicate. The spliced mRNAs were detected in lanes 5 and 7 (lower bands, open arrowheads) and an intron-retained mRNA was detected in lane 7 (upper band, closed arrowhead).

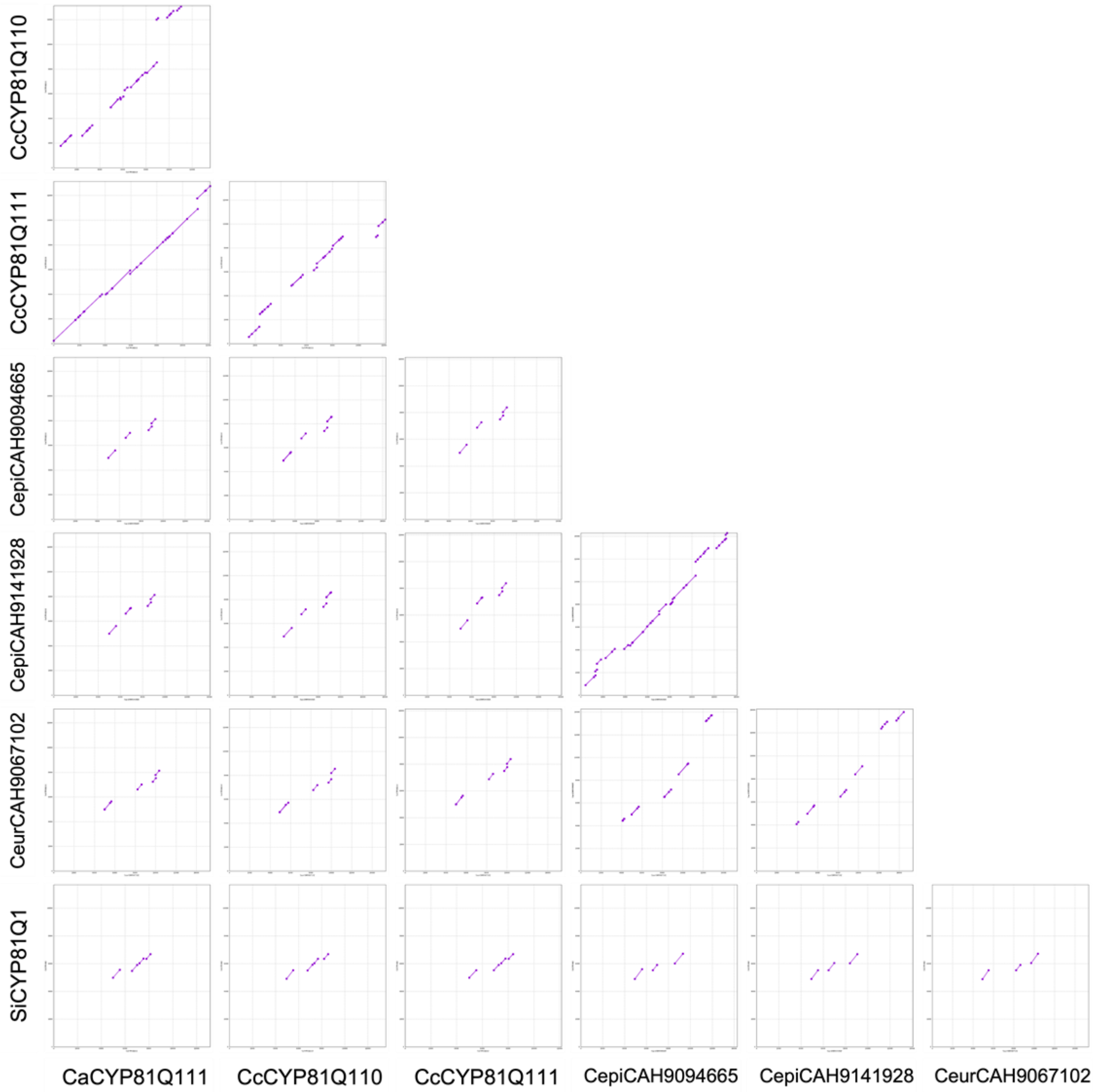

**Fig. S9.** Genome-by-genome alignment of genomic regions containing *CYP81Q*-related genes (plus/minus 5kbp)

Structural similarity in a broad genomic region (~10 kb with *CYP81Q* in the center) was evaluated using a dot-plotting tools. Alignment of *Cuscuta* genomic sequences performed using promer in the Mummer -v3.1.0 package (7). The plots show structural similarity of *C. campestris*, *C. australis*, *C. epithymum*, *C. europaea* and *S. indicum*. Purple dots indicate unique forward alignments. Structural similarities were visualized with dot plots using mummerplot. Only exons share structural similarity. These results are consistent with the idea that either 1) non-coding regions outside of *PSS* genes change easily due to loosened constraints for nucleic acid alteration as compared to the coding region, or 2) a small genomic fragment including the *PSS* ORF was transferred in a parasitism-mediated horizontal gene transfer event.

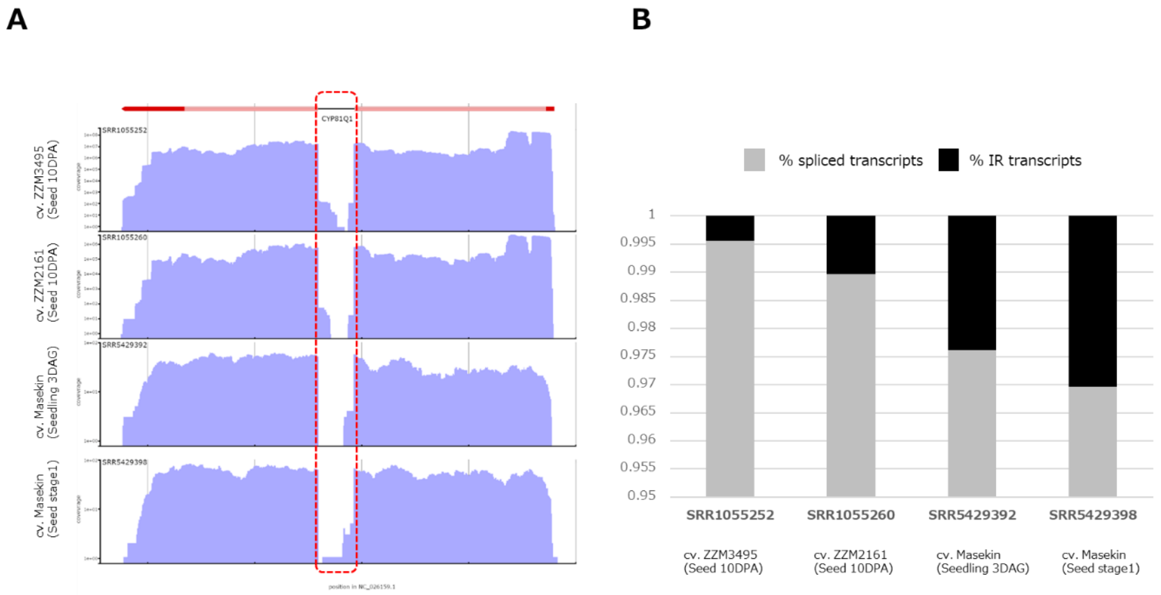

**Fig. S10.** Investigation of intron retention (IR) transcripts of *SiCYP81Q1* in sesame seeds

**A.** RNA-seq data from *Sesamum indicum* seeds of four cultivars (SRR1055252, SR1055260, SRR5429392, and SRR5429398) were mapped on the *S. indicum* genome using the *HISAT2* aligner (8). Identification of a small portion of RNA reads covering the intron of the *CYP81Q1* gene (NP_001306620), as indicated by the dotted red box, supports the idea that IR-transcripts of *CYP81Q1* gene are expressed as minor transcripts. **B.** Reads covering exon-intron junctions and reads in the intron are minor transcripts, accounting for < 3% of *CYP81Q1* transcripts. We counted numbers of reads (1) covering exon 1-exon 2, (2) mapping to exons, (3) covering exon 1-intron, (4) covering intron-exon 2, and (5) mapping to the intron, and then calculated splicing efficiency (6) using the following formula: (2)/{(2)+[(3)+(4)]}/2. Intron retention (7) was estimated as 1-(6). Relative abundance of spliced transcripts (6), black; of IR-transcripts (7), gray (graph on the right). These results suggest the presence of a very small portion of *IR-CYP81Q1* transcripts. Seed developmental stage was described in Ono et al., 2006 (9). DAG, days after gemination treatment; DPA, days post anthesis.

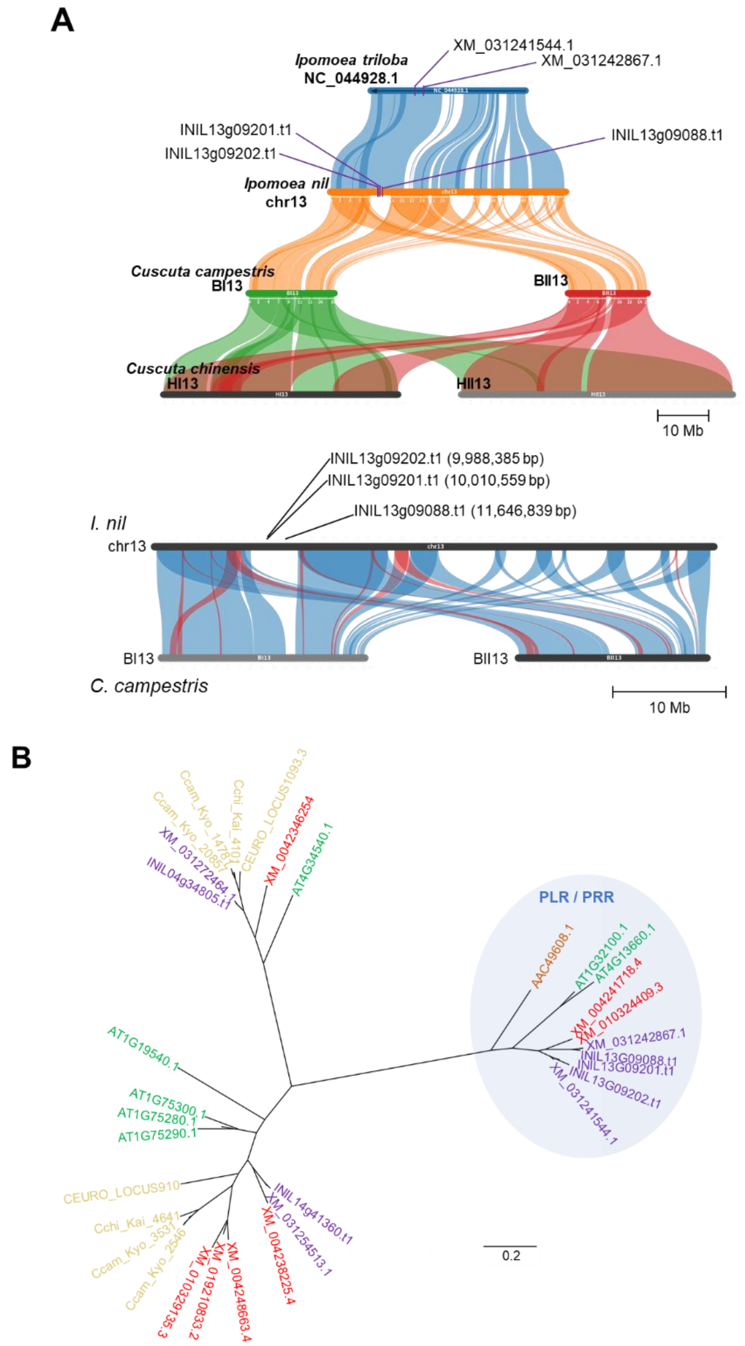

**Fig. S11.** Gene synteny and phylogenetic analyses of pinoresinol lariciresinol reductase (PLR)-related genes

**A.** Gene synteny comparison between PLR genes between *Cusucta* and *Ipomoea* genus. The genomic region containing Ipomoea *PLR*-like genes are lacked in two *Cuscuta* species although the genomic synteny of the most genes are highly conserved between the two phylogenetically related species. **B.** A phylogenetic tree of the *PLR*-related genes. Yellow, green, red, purple and orange color indicates the accession numbers of *PLR*-like genes derived from *Cuscuta* spp., *Arabidopsis thaliana*, *Solanum lycopersicum*, *Ipomoea* spp., and *Forysythia x intermedia*, respectively. No striking homologs of *Cuscuta* spp. was observed in the clade including FiPLR and *Arabidopsis* pinoresinol reductase (PrR) genes, suggesting that ancestral *Cuscuta* have lacked *PLR*-like at least before speciation intro *C. campestris* and *C. chinensis*.

C

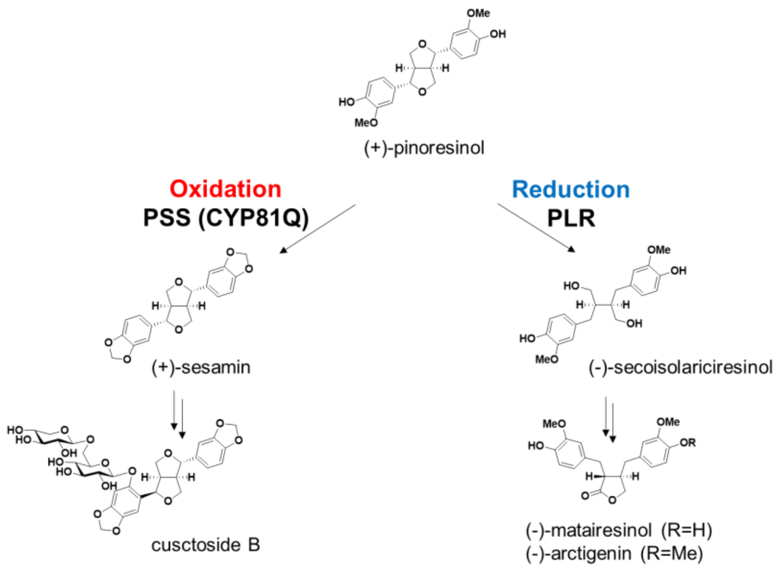

**Fig. S11.** Gene synteny and phylogenetic analyses of pinoresinol lariciresinol reductase (PLR)-related genes (continued)
**C.** Metabolic pathway of plant specialized lignans. Pinoresinol is the common substrate for PSS-catalyzed oxidation and PLR-catalyzed reduction. In *Cuscuta* spp., sesamin is further modified into cuscutoside through hydroxylation and glycosylation, while matairesinol and arctigenin are observed in *Ipomoea cairica*, suggesting functional *PLR* genes in the *Ipomoea* genome and metabolic competition between PLR and PSS.

**Dataset S1 (separate file).** Comparison of *CYP81Q*-related genes to *CcamCYP81Q111*

**Dataset S2 (separate file).** Nucleotide sequences of exon 2 of *CYP81Q*-related genes

**Dataset S3 (separate file).** Identifiers of proteins used in the phylogenetic analysis in Figure 2

**Dataset S4 (separate file).** Deduced amino acid sequences of *CYP81Q*-related genes

**Dataset S5 (separate file).** Genomic synteny among *Cuscuta* and related species

**Dataset S6 (separate file).** Estimated divergence times of *Cuscuta* and other plants

**Dataset S7 (separate file).** Transposons in the *CYP81Q*-related genes

**Dataset S8 (separate file).** Genome assembly containing Gene Feature Files used in the analysis of the number of introns per gene

**Dataset S9 (separate file).** Nucleotide sequences of the coding regions of the *CYP81Q*-related genes

**Dataset S10 (separate file).** Sequences of oligonucleotide primers and CDSs used in this study

### SI Methods

#### *Gene synteny and phylogenetic analyses of pinoresinol lariciresinol reductase (PLR)-related genes*

To identify homologs of *Forsythia* × *intermedia* PLR (AAC49608.1), DIAMOND blastp v2.1.12 (10) was used to search against amino acid sequences from *Cuscuta campestris* (t267557.G001, Plant GARDEN; https://plantgarden.jp/en/download/), *C. chinensis* (t132261, Plant GARDEN), *C. europaea* (GCA_945859875.1), *Ipomoea nil* (Asagao_chr_1.2, Morning Glory Genome Consortium; http://viewer.shigen.info/asagao/index.php), *I. triloba* (GCF_003576645.1), *Solanum lycopersicum* (GCF_000188115.5), and *Arabidopsis thaliana* (GCA_000001735.1). Homologous sequences identified were aligned using MAFFT v7.525 (--maxiterate 1000 –localpair) (11), and a phylogenetic tree was constructed using IQ-TREE v2.3.6 (-m MFP -bb 1000 -alrt 1000) (12). For synteny analysis, all genes located on the contig harboring the PLR homolog most closely related to *F.* × *intermedia* were subjected to homology search using DIAMOND blastp. Syntenic blocks were detected with MCScanX (13) and visualized using SynVisio (14) Bandi and Gutwin 2020).
